# Unfolding and identification of membrane proteins *in situ*

**DOI:** 10.1101/732933

**Authors:** Nicola Galvanetto, Zhongjie. Ye, Arin Marchesi, Simone Mortal, Sourav Maity, Alessandro Laio, Vincent Torre

## Abstract

Single-molecule force spectroscopy (SMFS) uses the cantilever tip of an AFM to apply a force able to unfold a single protein. The obtained force-distance curve encodes the unfolding pathway, and from its analysis it is possible to characterize the folded domains. SMFS has been mostly used to study the unfolding of purified proteins, in solution or reconstituted in a lipid bilayer. Here, we describe a pipeline for analyzing membrane proteins based on SMFS, that involves the isolation of the plasma membrane of single cells and the harvesting of force-distance curves directly from it. We characterized and identified the embedded membrane proteins combining, within a Bayesian framework, the information of the shape of the obtained curves, with the information from Mass Spectrometry and proteomic databases. The pipeline was tested with purified/reconstituted proteins and applied to five cell types where we classified the unfolding of their most abundant membrane proteins. We validated our pipeline by overexpressing 4 constructs, and this allowed us to gather structural insights of the identified proteins, revealing variable elements in the loop regions. Our results set the basis for the investigation of the unfolding of membrane proteins in situ, and for performing proteomics from a membrane fragment.

## Introduction

Mapping and recovering the structure of membrane proteins in their native environment, *i.e.* embedded in a native cellular membrane and at room temperature is a challenging aim: indeed the most successful tool for this purpose is cryo electron microscopy, but it requires frozen samples, and precise single particle measurements can be achieved only with purification.

In fact, much of what we know about the mechanics and the structure at room temperature of cell membranes, of polymers (Carrion-Vazquez et al., 1999; Sarkar et al., 2005) and of membrane proteins (Al-Rekabi and Contera, 2018; Casuso et al., 2012; García-Sáez et al., 2007) comes from atomic force microscopy (AFM) which can operate in liquid environment. Single Molecule Force Spectroscopy (SMSF) uses the tip of an AFM to apply a stretching force to unfold a single protein. SMFS provides a 1D force-distance (F-D) curve which encodes the unfolding pathway so that from the analysis of the sequence of force peaks it is possible to identify the folded domains and their variability (Engel and Gaub, 2008). It has been recently confirmed that from the 1D amino-acid sequence of a protein it is possible to accurately determine its 3D structure (Jumper et al., 2021), therefore it is tempting to continue exploring which information can be recovered from the 1D F-D curves provided by SMFS.

Hitherto SMFS has been mostly used to study the unfolding mechanics of purified proteins, in solution or reconstituted in lipid bilayers. The information that is possible to obtain (e.g. mechanical stability(Sumbul et al., 2018a; Thoma et al., 2015), structural heterogeneity(Hinczewski et al., 2016)) depends on the physical and chemical properties of the cell membrane(Maity et al., 2015; Thoma et al., 2018). However, native membranes are extremely complex, hosting hundreds of different proteins in an environment which is highly heterogeneous even at the single-cell level, and typically different in different cells.

In this manuscript we aim to bridge the gap between all these recent breakthroughs by attempting to identify and recover structural information of membrane proteins embedded in biological samples, i.e. in their native environment and at room temperature.

We describe a complete pipeline, including the experimental methods and the data analysis, which represent a first step forward in this direction. The pipeline allows to identify membrane proteins obtained from SMFS on single cells native membranes. Firstly, we developed a technique to extract a piece of the membrane suitable for SMFS so to obtain millions of F-d curves from native biological membranes. Secondly, we used an unsupervised clustering procedure to detect clusters of similar unfolding curves. Finally, we implemented a Bayesian meta-analysis using information from Mass Spectrometry and protein structure databases that allows to identify a limited list of candidates of the unfolded proteins.

We first characterized the unfolding of purified membrane proteins reconstituted *in vitro* in lipid bilayers, then we focused on the native cell membranes of hippocampal neurons, dorsal root ganglia (DRG) neurons, neuroblastoma cells and the plasma and disc membrane of rod outer segments. Our protocol — even if it cannot compete with the identification accuracy of mass spectroscopy performed on thousands of cells, or with the data quality of a SMFS experiment performed on a single purified protein in an artificial membrane — accomplishes a different task, and in each of these cases it allows gathering biologically meaningful information that cannot be captured otherwise, even at the single cell level.

## Results

### Unfolding proteins from isolated cell membranes

The first ingredient of our pipeline is an unroofing method(Galvanetto, 2018) to isolate the apical part of single cell membranes containing membrane proteins with negligible contamination of cytoplasmic proteins (Fig. S1–S3). We sandwiched a single cell between two glass plates, the culture coverslip and another plate mounted on the AFM itself (see Fig. 1B-C, triangular coverslip). The triangular coverslip is coated with polylysine which favors membrane adhesion. A rapid separation of the plates, permits the isolation of the apical membrane of the cell (see Fig. 1D-E). The method is reliable (n=105, ~80% success rate) with cell types grown on coverslips. With non-adherent cells, like freshly isolated rods, membranes were isolated with a lateral flux of medium(Clarke et al., 1975) (see Methods). During the unroofing process, we verified the absence of any adsorption of cytoplasmic material like tubulin, actin, mitochondria or free cytoplasmic proteins from the unroofed membrane patch where only the membrane proteins that are hold by the lipids are present (see Fig. S2–S4).

**Fig. 1.**
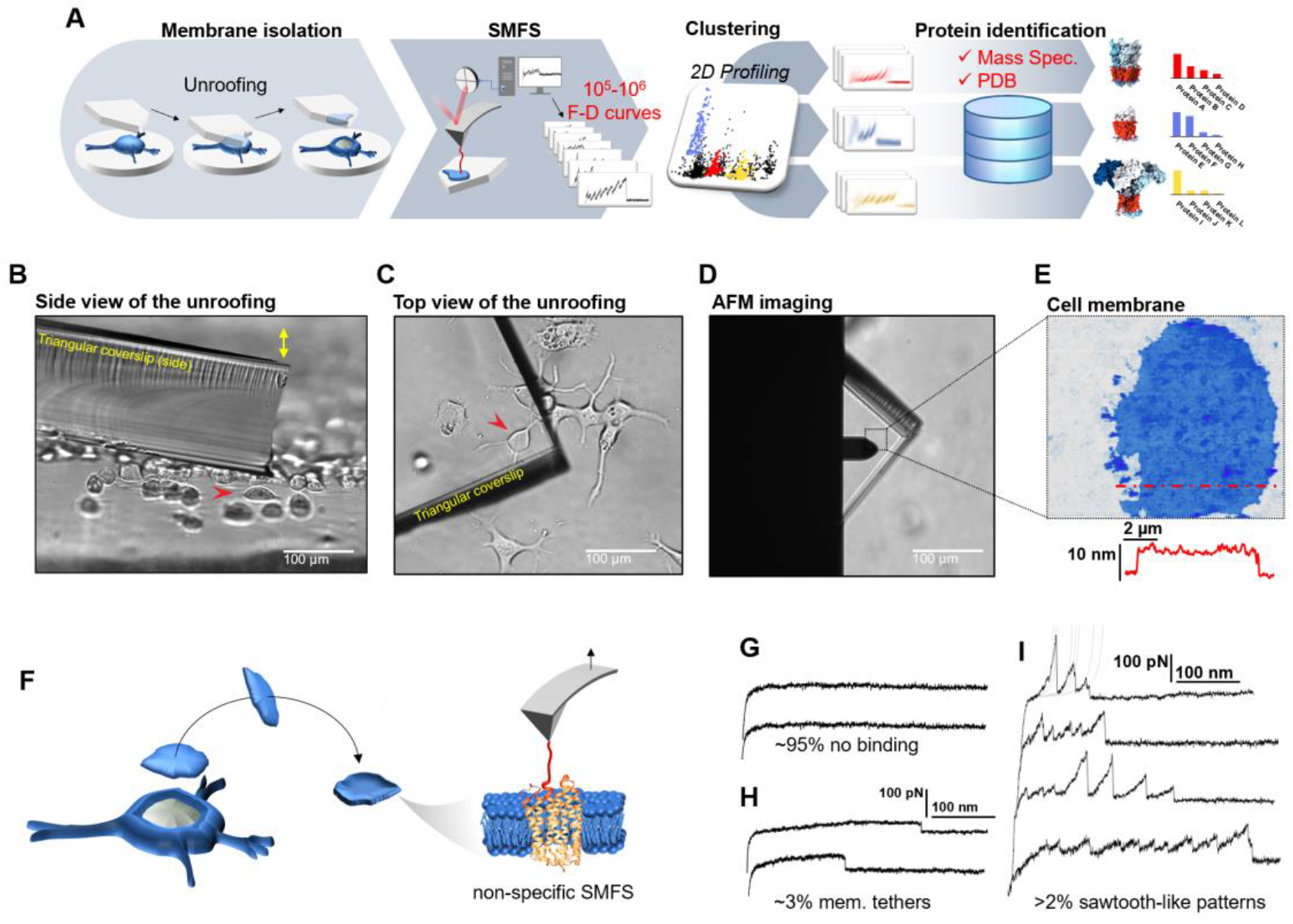
Experimental method for single-cell membrane isolation and protein unfolding. (**A**), workflow of the method in four steps: isolation of the apical membrane of single cells; AFM-based protein unfolding of native membrane proteins; identification of the persistent patterns of unfolding and characterization of the population of unfolding curves; Bayesian protein identification thanks to mass spectrometry data, Uniprot and PDB. (**B**), side view and (**C**), top view of the cell culture and the triangular coverslip approaching the target cell (red arrow) to be unroofed. (**D**), positioning of the AFM tip in the region of unroofing. (**E**), AFM topography of the isolated cell membrane with profile. (**F**), cartoon of the process that leads to SMFS on native membranes. Examples of F-D curves of (**G**), no binding events; (**H**), constant viscous force produced by membrane tethers during retraction; (**I**), sawtooth-like patterns, typical sign of the unfolding of a membrane protein.

We verified with the AFM (Fig. 1F) that the isolated membrane patches have a height of 5-8 nm with roughness in the order of 1 nm. Then, we performed conventional SMFS(Oesterhelt et al., 2000) collecting more than two millions of F-D traces from all the samples (hippocampal neurons, DRG neurons, neuroblastoma, rods and rod discs, see below). Among the obtained curves, ~95% shows no binding (Fig. 1G), ~3% shows plateau ascribable to membrane tethers (Fig. 1H), while the remaining ~2% displays the sawtooth-like shape that characterizes the unfolding of proteins(Oesterhelt et al., 2000) (Fig. 1I) with each tooth fitting the worm-like chain (WLC) model with a persistence length of ~0.4nm (Fig. S5)(Li et al., 2002). These ~50,000 traces are dramatically heterogeneous, differing in total length, number of peaks, maximum force, etc. This is not surprising, since cell membranes contain a large number of different proteins, each hosted in different local environment. In the following we describe a procedure which allows, to some extent, disentangling this bundle.

In SMFS it is assumed that the binding between the cantilever and the protein occurs either at the C- or at the N-terminus, and that the protein is fully unfolded by the tip. However, some traces suggest that also other events take place: i) the simultaneous attachment of two or more proteins to the tip(Walder et al., 2017), ii) the incomplete unfolding of the attached protein(Tanuj Sapra et al., 2006), iii) the binding of the AFM tip to a loop of the protein instead of to a terminus end (Fig. S6, C to F).

#### i) *Attachment of multiple proteins* (Fig. S6D)

the resulting F-D curves will not have a recurrent pattern; if two proteins form a complex, the resulting spectrum is the sum of the two individual spectra: that causes deviations of the measured persistence length (Fig. S7). The simultaneous unfolding of multiple proteins is also characterized by force changes and varying persistence length (Fig. S6D, G and S7).

##### ii) *Incomplete unfolding of the protein* (Fig. S6E)

if the tip prematurely detaches from the terminus, the F-D curve has similar but shorter pattern compared to a complete unfolding (Fig. S6C). The fraction of curves that prematurely detaches has been reported to be ~23% of the fully unfolded protein(Tanuj Sapra et al., 2006).

##### iii) *Binding of the AFM tip to a loop* (Fig. S6F)

this case is equivalent to the attachment of multiple proteins. However, if the attachment of the cantilever tip to a loop occurs with some consistency, we will obtain a recurrent pattern with the features described in case i) (deviation of persistence length during intersection, 2 major levels of unfolding force, see Fig. S7).

Therefore, there is no doubt that a large fraction of F-D curves will not represent the proper unfolding of a single protein. But inspired by the successes of single particle Cryo-EM, which can produce atomic-resolution structures, despite selecting less than 20% of the protein images(Yi et al., 2019), we attempted a similar approach on SMFS data, under the hypothesis that ‘bad’ events are likely to produce *non-recurrent* patterns of unfolding.

The key tool to recognize recurrent patterns is an automated clustering method developed in our group(Ilieva et al., n.d.) based on density peak clustering(Rodriguez and Laio, 2014) (Fig. S11, see Supplementary Methods for a detailed description of the algorithm). In short, this approach detects statistically dense F-D patterns occurring often in the sample (clusters). It is based on a previously established ‘similarity distance’(Marsico et al., 2007) in the context of SMFS and, remarkably, it does not require to pre-set neither the shape nor the number of clusters.

### Benchmarking the analysis with mixtures of purified proteins reconstituted in lipid bilayers

We first tested the consistency of our pipeline with three known membrane proteins. We reconstituted in lipid bilayers(Shen et al., 2013) highly purified Channelrhodopsin (ChR1)(Nagel et al., 2002), Bacteriorhodopsin (bR)(Oesterhelt et al., 2000) and the cyclic adenosine monophosphate (cAMP) gated channel SthK(Marchesi et al., 2018) (Fig. 2A), and we performed SMFS experiments. bR, ChR1 and SthK were reconstituted *in vitro* one at a time. Since we did not have control of the protein orientation in the bilayer, we expected 2 main F-D patterns, and therefore two clusters for each sample: one representing the unfolding of the protein from the N-terminus and the other from the C-terminus. The clustering algorithm identified 2 almost indistinguishable clusters for bR ( see Supplementary Methods), and the result is in agreement with the fact that the unfolding of bR from the two termini is almost symmetric as previously reported (Kessler and Gaub, 2006) (Fig. 2D). In the case of ChR1 we obtained 3 clusters where the second cluster — cluster C2 — represents the partial unfolding of ChR1 from the C terminus because the peaks of C2 match perfectly the first three peaks of the full unfolding (cluster C1), while cluster C3 perfectly match the unfolding from the N terminus (Fig. 2B). Sthk generated 2 clusters, cluster S1 and S2 (Fig. 2C), as expected (all the clusters are shown in Fig. S8). For all the three proteins, we observed that: (i) the value of Lc_max_ — i.e. the estimated length of the segment of a.a. stretched in our SMFS experiments — is within 5% equal to the total length of the protein minus the free length of the N-terminus (C-terminus) domain if the cantilever tip attached to the C-terminus (N-terminus) (see Fig. 2B, C and Fig. S8B); and that (ii) the force peaks of the clusters of F-D curves colocalize with the unfolded regions or loops of their structures (see Fig. S8, A and C).

**Fig. 2.**
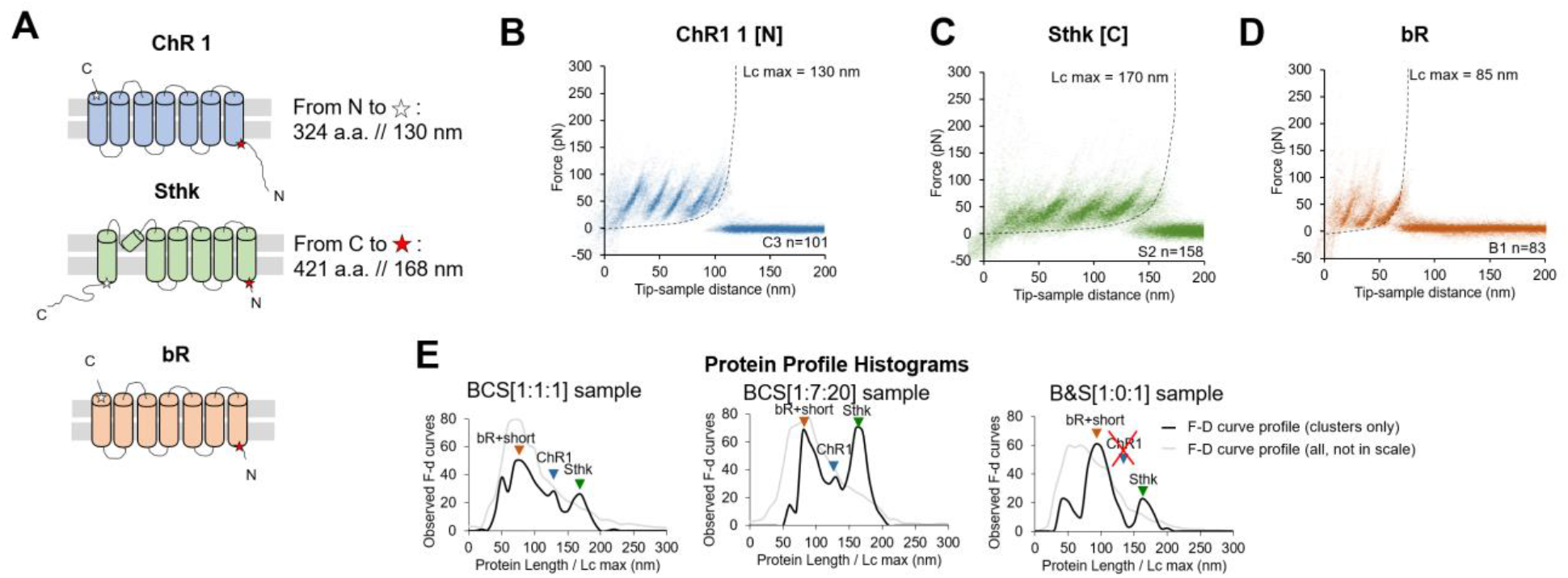
Unfolding of reconstituted mixtures of membrane proteins. (**A**): scheme of the structure of the three purified proteins used in the in vitro preliminary step: ChR1, Sthk and bR (cylinders represent alpha helices). (**B**): superimposition of 101 unfolding curves (density plot) of the full unfolding of ChR1 from the N terminus. **(C)**: density plot of the full unfolding of Sthk from the C terminus.. (**D**): density plot of the full unfolding of bR. (**E**): protein profile, i.e. the histogram of the maximal contour length (Lc max) of all the F-D curves in the clusters (black line), and of all the F-D curves collected from the sample (grey line, not in scale), from samples with mixtures of bR, ChR1 and Sthk with a relative abundance of 1:1:1, 1:7:20 and 1:0:1 (bR-Sthk only). Arrows indicate where the F-D curves of (**B**), (**C**) and (**D**) accumulate in the histogram. The peak at ~90 nm indicated by bR+short also contains the shorter clusters of ChR1 and SthK shown in Fig. S8, that also appear in the mixtures.

Next, we simultaneously reconstituted two proteins in lipid bilayers (see Methods) obtaining the same clusters associated with the unfolding patterns observed in the single protein reconstitution (Fig. S9). We finally reconstructed all the three proteins together, with different relative abundances: 1:1:1 and 1:7:20 respectively for bR, ChR1 and SthK. We observed that the number of traces per cluster scaled approximately with the abundance of the preparation (Fig. S10).

The histogram of the maximal contour length (Lc max) of all the F-D curves (Figure 2E) encodes the protein content of the sample and allows a rapid detection of the presence and of the abundance of a given protein in sub-femtogram samples, providing also a qualitative indication on the concentration of a given protein. The success of this preliminary *in vitro* investigation prompted us to move to native membranes.

### Clustering SMFS data from native membrane proteins

Next, we applied our pipeline to DRG, hippocampal neurons, neuroblastoma, rod outer segments and rod discs membranes (Fig. 3A-E) and we obtained 301,654 curves from the hippocampal neurons, 413,468 from DRG neurons, 623,482 from neuroblastoma, 386,128 from rods and 221,565 from rod discs. From these F-D traces we applied our clustering procedure that identified 15, 10, 11, 8 and 5 statistically dense pattern of unfolding present in the datasets (Fig. 3F-J). These traces show that clean unfolding patterns can be detected even in patches of native membranes.

**Fig. 3.**
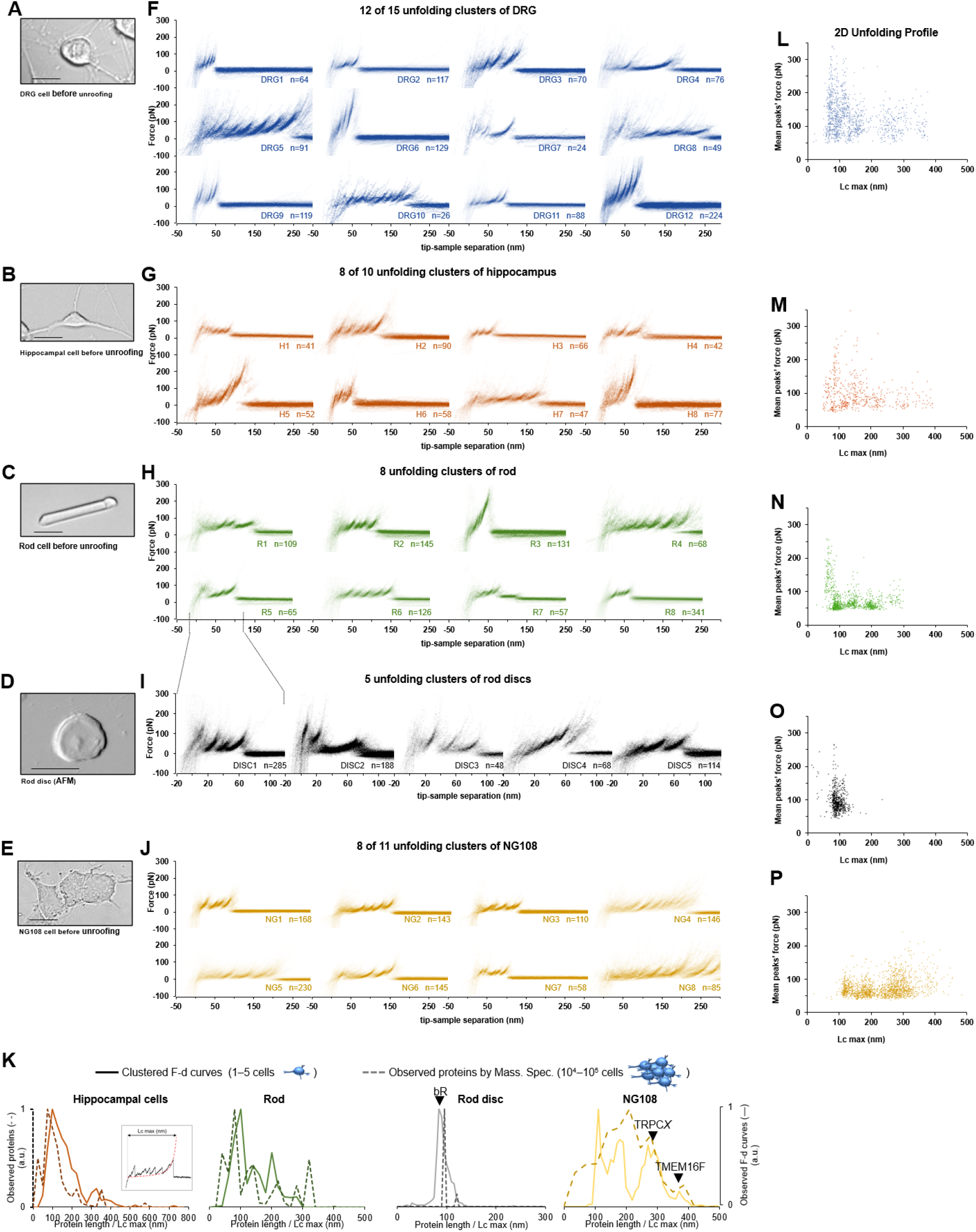
Unfolding clusters in native cell membranes. Bright field image of (**A**): dorsal root ganglia neuron; (**B**): hippocampal neuron; (**C**) rod before unroofing (scale bar 15 μm). (**D**): AFM error image of an isolated disc (scale bar 1 μm), (**E**): NG 108 cells. (**F, G, H, I, J**): examples of obtained clusters from the native membranes shown as density plots, that is superposition of the n unfolding curves agglomerated and colored as a heat map. (**K**): comparison of the Protein profiles obtained with mass spectrometry vs. the F-D curves obtained with SMFS shows a good correlation. The protein profile from mass spectrometry is the normalized histogram of the total number of proteins found in the sample, considering their abundance. The protein profile from SMFS is the histogram of the maximal contour length (Lc max) of all the F-D curves selected with our clustering procedure. (**L, M, N, O, P**): representation of all the clustered F-D curves in 2D: x-axis is the maximum contour length; y-axis is the average unfolding force (DRG: n = 1255; hippocampus: n = 563; rod: n = 1039; disc: n = 703, NG108: n = 1591).

We identified four major classes of clusters: *Short curves with increasing forces:* clusters DRG12, H5, H8 and R3 shows repeated peaks (ΔLc 10-20 nm, distance between consecutive peaks) of increasing force up to 400 pN. *Long and periodic curves:* R6, H7 and DRG10 display periodic peaks of ~100 pN and with a ΔLc of 30-40 nm whose unfolding patterns are similar to what seen when unfolding LacY(Serdiuk et al., 2016). *Conventional short curves:* the majority of the identified clusters like DRG1, H3, R8 and all clusters from the rod discs have F-D curves with a total length less than 120 nm with constant or descending force peaks. These F-D curves share various features with the opsin family unfolded in purified conditions(Engel and Gaub, 2008), *e.g.* a conserved unfolding peak at the beginning (at contour length < 20 nm) associated to the initiation of the denaturation of the protein. We found also *unconventional* clusters such as DRG7, DRG8 and R7: DRG8 exhibits initial high forces and with variable peaks followed by more periodic low forces; while cluster R7 has a conserved flat plateau at the end of the curve of unknown origin.

A compact representation of clustered F-D curves becomes more important in native samples where the information stored in the SMFS data is more complex (see Fig. 3L-P). The cancerous NG108-15 cells have few short and not particularly stable membrane proteins compared to neural cells which instead have many clusters of proteins under 100 nm that unfold even above 200 pN. On the contrary, NG cells have a higher fraction of long and stable proteins than neurons.

Furthermore, in native membranes we can directly compare the expected protein profile obtained from mass spectrometry (usually obtained from thousands of cells, see Methods) with the F-D curves profile of just few cells (Fig. 3K) which shows a good agreement in particular in the region above 100 nm of protein length. These results suggest that protein length detection of SMFS — restricted on clustered traces according to our protocol — provides a semi-quantitative proxy for mass detection at the single cell membrane level.

### Molecular identification of detected clusters

Having identified clusters of F-D curves from native membranes that approximates the overall population of membrane proteins (Fig. 3K), the next question is: *can we identify the membrane proteins whose unfolding corresponds to the identified clusters (Fig. 3)?*

For this purpose, we developed a Bayesian method providing candidate proteins on the basis of the information present in the data from Mass Spectrometry (ProteomeXchange) and in other proteomic databases (Uniprot, PDB) — combined with the empirical results obtained from SMFS literature(Bosshart et al., 2012; Cisneros et al., 2005; Ge et al., 2016; Kawamura et al., 2013; Kedrov et al., 2004; Klyszejko et al., 2008; Maity et al., 2015; Möller et al., 2003; Oesterhelt et al., 2000; Sapra et al., 2009; Serdiuk et al., 2016; Thoma et al., 2017, 2012). The Bayesian identification (Fig. 4A) is based on two steps: firstly, the crossing of information between the clusters with the results of Mass Spectrometry analysis of the sample under investigation (hippocampal neurons, discs, etc.); secondly, a refinement of the preliminary candidates using additional information (structural and topological) present in the PDB and Uniprot databases. The first step relies on the comparison of Lcmax of each cluster with the length of proteins identified in the Mass Spectrometry data (Chen et al., 2006; Kwok et al., 2008; Panfoli et al., 2008). By using the protein abundance included in the Mass Spectrometry data as the prior, a first list of candidate proteins is obtained as well as their probabilities (Fig. 4A III). The SMFS-literature contains 14 examples of unfolded membrane proteins allowing a comparison between the Lcmax of the measured F-D curves and the real length of the same protein completely stretched (Fig. 4B). Therefore, we extrapolated the first conditional probability of our Bayesian inference, showing that on average, the value of Lcmax corresponds to 89% of the real length of the protein (R^2^=0.98). When the structure of candidate proteins is known (helices, sheets), these probabilities are further refined considering e.g. the force of unfolding or the position of the loops which are known to constrain the possible unfolding patterns (Fig. 4C-F, see supplementary methods for the formal description).

**Fig. 4:**
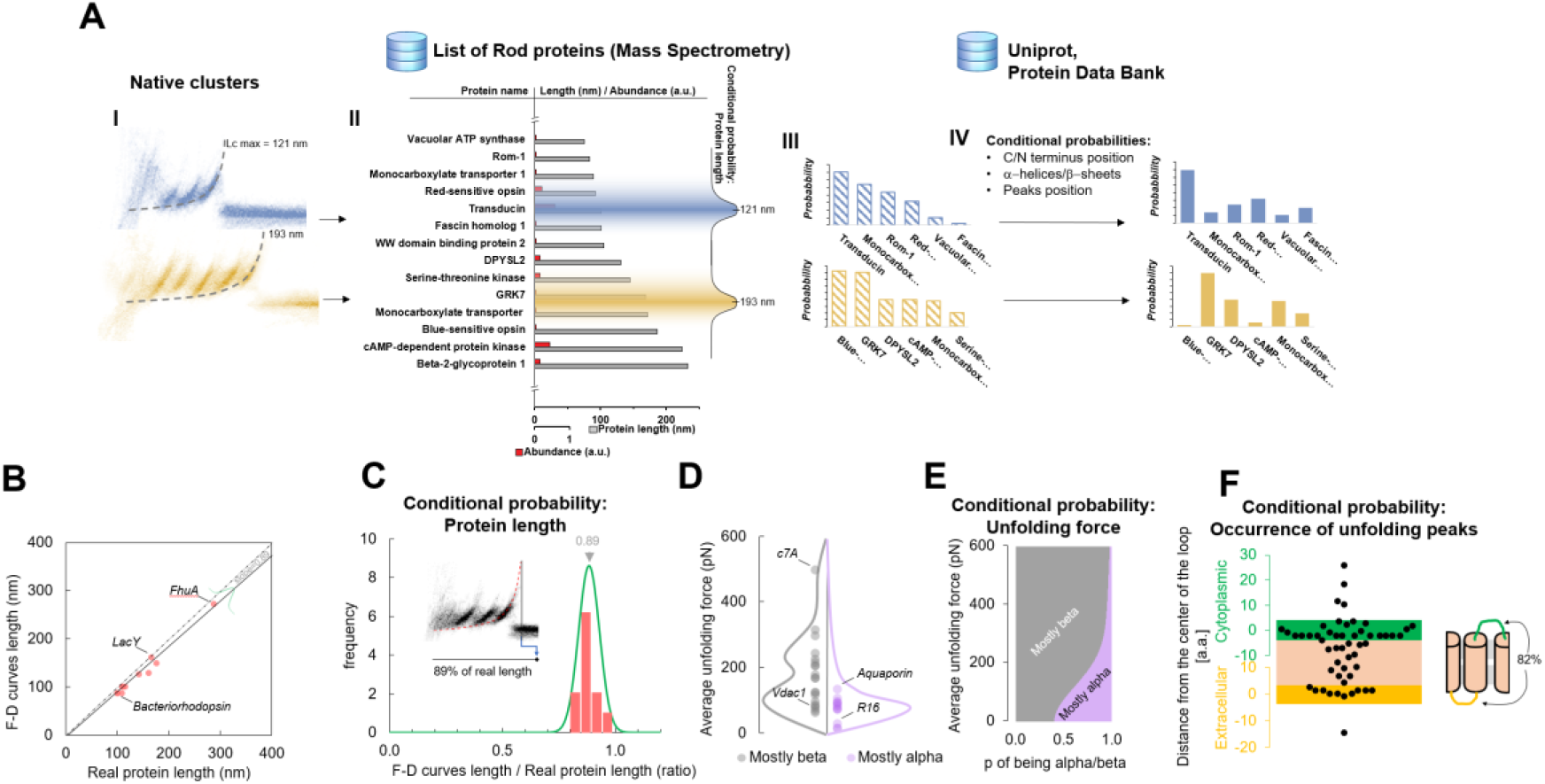
Bayesian identification and conditional probabilities. **A**, logical workflow of the Bayesian steps: selection due to total length and abundance (mass spectrometry), refinement with structural and topological information (PDB and Uniprot). **B**, Comparison of the real length of the protein vs. the measured maximal contour length of the F-D curves in 14 SMFS experiments on membrane proteins. **C**, Conditional probability of the observed maximal length of the clusters obtained from **B**. **D**, Comparison of the force necessary to unfold beta sheets and alpha helices in 22 SMFS experiments. **e**, Conditional probability of the observed unfolding forces obtained from **D**. **F**, Conditional probability for the occurrence of unfolding peaks extracted from SMFS literature (see methods). Unfolding peaks occurs most likely in the loops (82%) than in transmembrane domains (n_peaks=54, from 11 SMFS experiments of different membrane proteins). The points in the green (yellow) region represents unfolding peaks occurred in a cytoplasmic (extracellular) loop, the point in the pink area occurred in a transmembrane domain. The points above the green and below the yellow regions occurred in cytoplasmic and extracellular domains, respectively. The scale is approximate because in rare occasion loops are longer than 10 nm.

Like in Mass Spectrometry, this identification is probabilistic (Fig. 5), and it allows to reduce the number of possible candidates from hundreds to 1 to 4 proteins.

**Fig. 5.**
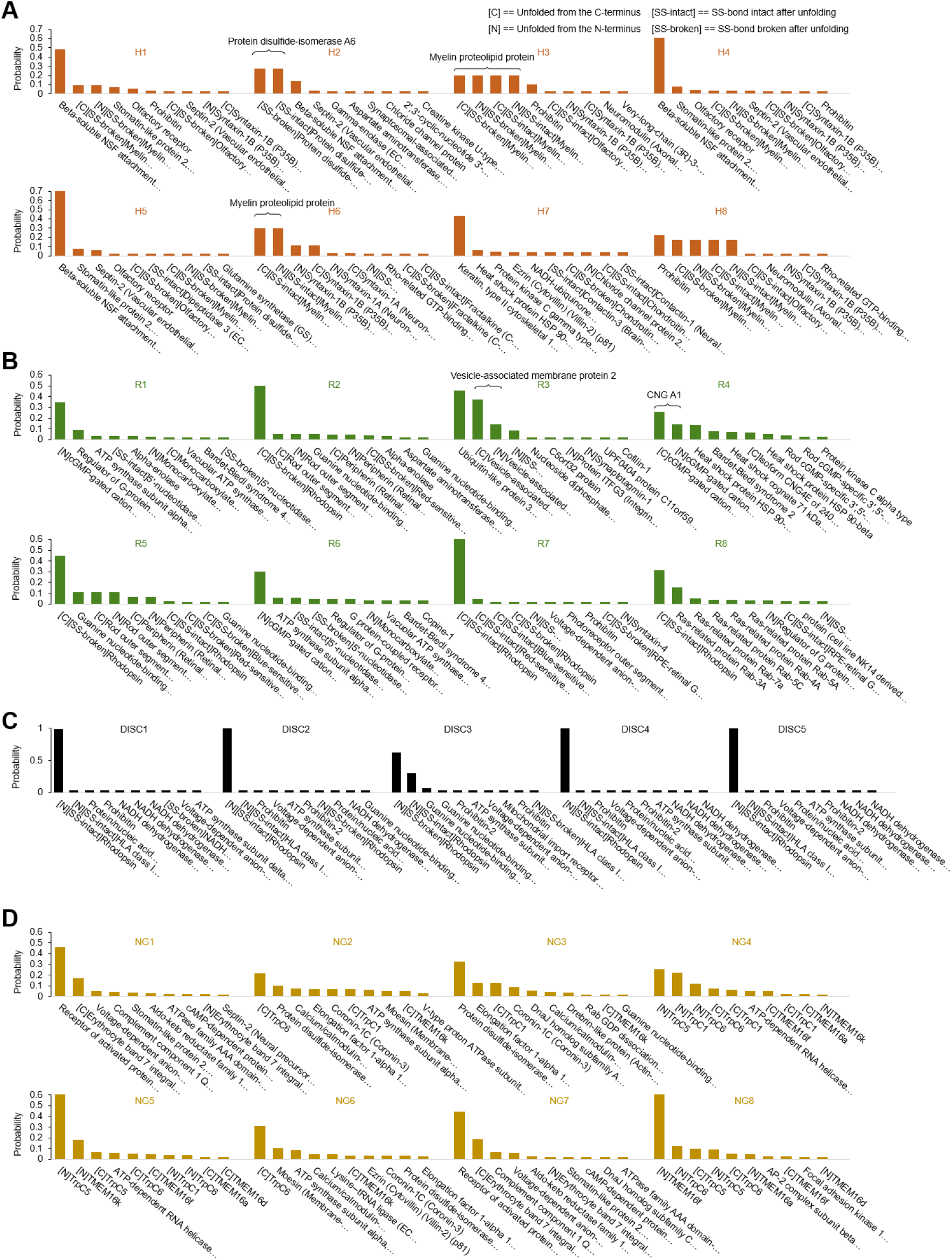
Bayesian identification of the unfolding clusters. Most probable candidates for the unfolding clusters found in a, hippocampal neurons; b, rods; c, rod discs; d, NG108-15 cells.

To prove the reliability of the proposed pipeline protocol, we focused on 4 clusters found in NG108-15 cells where the Bayesian method assigned a high probability to the membrane proteins TMEM16F, TRPC1, TRPC5 and TRPC6. Reverse Transcription Polymerase Chain Reaction and Western Blot analysis (Wu et al., 2007) (Fig. S17B) confirm that these membrane proteins are abundantly expressed in neuroblastoma NG108-15 cells. We overexpressed the construct 6xHis-N2B-*protein*-GFP, where *protein* is the molecular candidate to be validated (Fig. 6A–B, Table S1). The N2B is a chain of 204 aa which can be unfolded with a force less than 10 pN providing a well-known signature(Linke and Grützner, 2008), and GFP is the Green Fluorescent Protein used to identify which NG108-15 cell was successfully transfected, and if the proteins were present in the unroofed membrane patch (Fig. 6C).

**Fig. 6.**
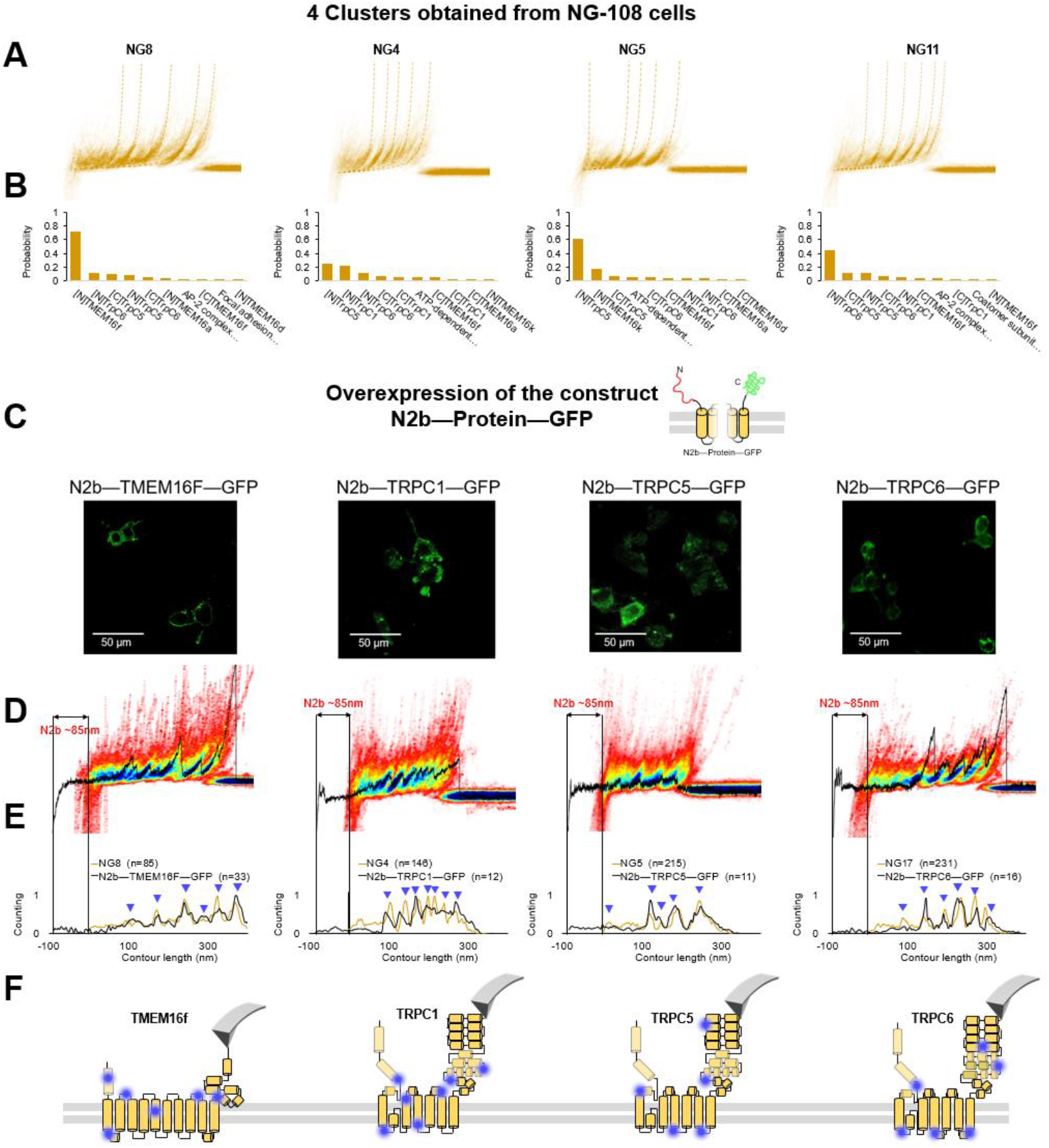
Validation of the method with a comparison between native clusters and those obtained with overexpression of a construct with the N2B signature at the N-terminal and GFP at the C-terminal. **A**: four clusters found in the dataset of F-D traces pulled from NG108-15 cells wild type. **B**: the Bayesian identification of the clusters in **A** with the candidate membrane proteins. **C**: confocal images of NG108-15 cells overexpressing the construct N2b—Protein—GFP, as reported by the green fluorescence emitted by GFP. **D**: from left to right, color density plots (blue indicating a colocalization among F-D traces of more than 80 %) of the clusters superimposed to a F-D curve bearing the N2B signature (85 nm segment with a flat force below 10 pN) obtained in the sample overexpressing the candidate protein TMEM16F, TRPC1, TRPC5 and TRPC6, respectively. **E**: Global histogram of Lc of the clusters in **A** and the clusters with the N2B signature from cells transfected with the corresponding construct **F**: position of the most likely rupture regions of the proteins according to **E**. superimposed to the cartoon of the Cryo-EM structure available in the PDB.

We compared the clusters obtained without overexpression (NG8, NG4, NG5 and NG11) with the new clusters obtained from the unfolding of the constructs N2B-TMEM16F-GFP, N2B-TRPC1-GFP, N2B-TRCP5-GFP and N2B-TRPC6-GFP from the transfected cells (Fig. 6D, E). These F-D traces had the well-known signature of the N2B domain consisting in a flat portion with a length of ~85 nm, which was followed by force peaks almost exactly matching those observed in the corresponding density plots (Fig. 6D). This matching was also confirmed by the correspondence of the cumulative peaks observed in the global histogram of the contour length of the NG8, NG4, NG5 and NG11 clusters with those obtained from neuroblastoma cells transfected with the constructs N2B-TMEM16F-GFP, N2B-TRPC1-GFP, N2B-TRCP5-GFP and N2B-TRPC6-GFP translated of ~85 nm because of the presence of the N2B (Fig. 6E).

This pipeline demonstrates that from the obtained clusters (Fig. 3) it is possible to derive the unfolding spectra of membrane proteins in their almost-native conditions without the need of purification. The superposition of the peaks (Fig. 6D–E) confirms the identification and allows to map the unfolding positions along the tertiary structure of the proteins (Fig. 6F).

### Structural insights from SMFS

The confirmation of the molecular identity of the unfolded proteins allows a better understanding of their molecular structures obtained from Cryo-EM, in particular of the not-well folded regions. SMFS detects the unfolded domains of a protein and can characterize their variability where the more powerful Cryo-EM precisely determines the position of the atoms of the well folded domains. The unfolding of the construct N2B-TRPC6-GFP as well as the endogenous TRPC6 show force peaks at 143, 193 and 230 nm with different properties (Fig. 7). The peak at 143 nm is always well defined, but the force peaks at 193 nm and at 230 nm — occurring at the extracellular loop E1 between S1 and S2 and at the extracellular loop E2 between S3 and S4, respectively (see positions in Fig. 7B) — show variability in the location and strength of the secondary peaks (Fig. 7D and E). The unfolding behavior observed at 193 nm and 230 nm reflects the presence of small structural elements(Yu et al., 2017) which are present in some but not all instances (see inserts in Fig. 7D-E). The peak at 143 nm, in contrast shows a clear and reproducible unfolding behavior (Fig. 7C) typical of a defined and well folded structure(Rico et al., 2013; Takahashi et al., 2018).

**Fig. 7.**
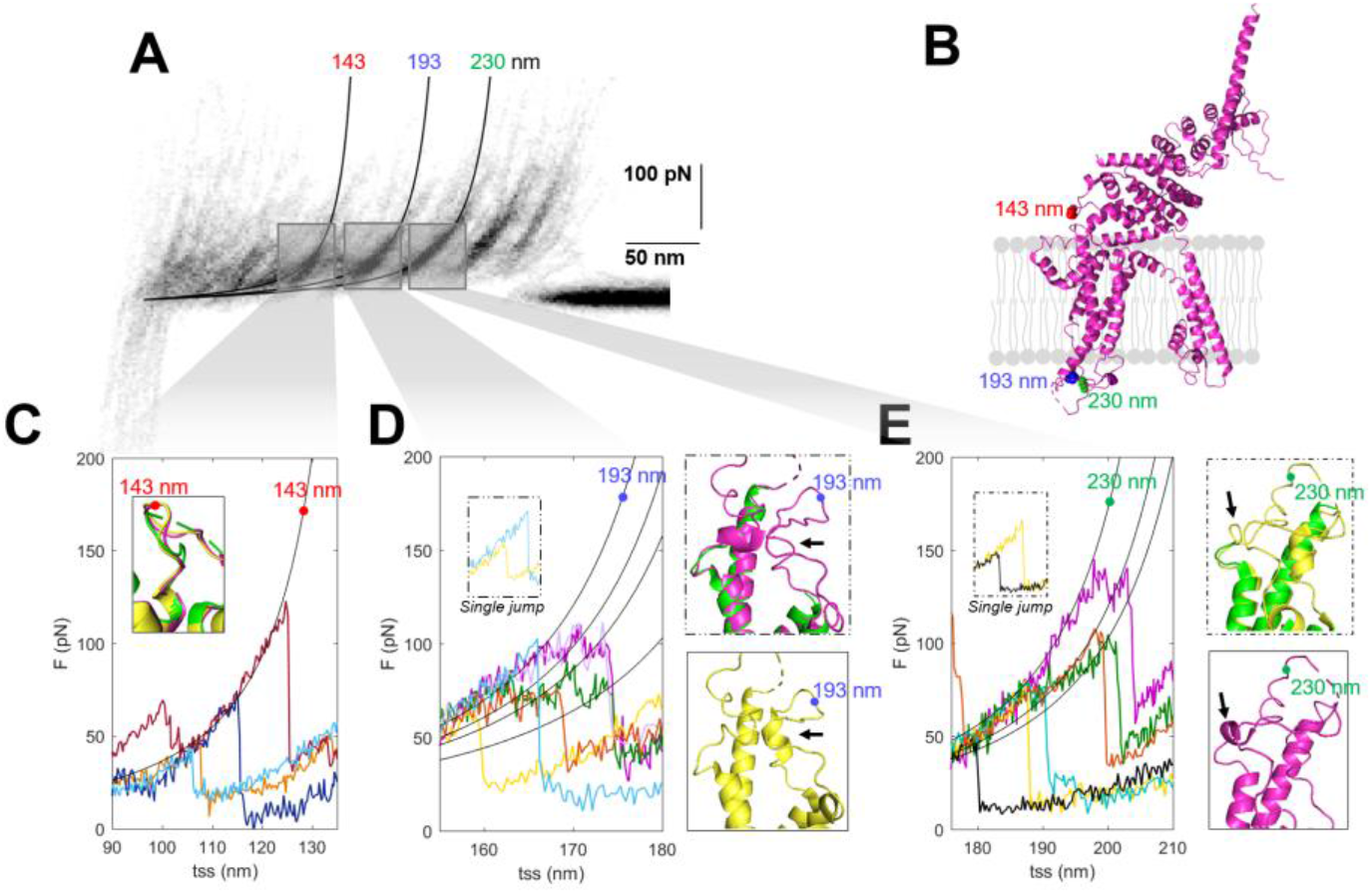
Structural segmentation in the loops of TRPC6. **A**: Density plot of F-D curves of TRPC6 (validated from cluster NG11). **B**: 3D structure of TRPC6 (PDB: 6UZA) with highlighted the position corresponding to the unfolding peaks in A. **C**: Representative F-D curves showing the single peak at 143 nm. Insert: comparison of the agreement with the three proposed structures of TRPC6 (PDB ID, green: 5YX9; purple: 6UZA; yellow: 6UZ8). **D**: Representative F-D curves showing the variable behavior at 193 nm where ~20% of the F-D curves shows a single jump that suggest no structural fragmentation of the loop like in the green/purple structures. Right inserts: comparison of the structure of the loop obtained by Cryo-EM where only the yellow structure show the presence of a 2-turn helix. **E**: Representative F-D curves at 230 nm where ~30% of the F-D curves shows a single jump that suggest no structural fragmentation of the loop like in the green/yellow structures. Right inserts: comparison of the structure of the loop obtained by Cryo-EM where only the purple structure show the presence of a 1-turn helix.

In the light of these results, we analyzed the available Cryo-EM structures of TRPC6. There are three published structures (Bai et al., 2020, p. 6; Tang et al., 2018, p. 6): these three Cryo-EM structures are very similar — and almost identical — particularly in the long transmembrane α-helices, but differ in the connecting loops where short α-helices are present only in some of these structures (Fig. 7 C-E, inserts). These short α helices in the loops E1 and E2 are the small structural elements at the origin of the segmentation observed in the F-D traces (Fig. 7 D-E). However, as shown in the structures in yellow, the α helices are not always observed as the single-peak F-D curves suggest. Therefore the most likely conclusion from our SMFS data is that, at room temperature and in their native environment, the different Cryo-EM structures coexist and the connecting loops E1 and E2 have a variable structure.

The observed variability is not restricted only to loop E1 and E2 of TRPC6: also the loop pre-S1 and S1 of TRPC5 shows a similarly complex unfolding behavior as opposed to the unfolding of the loop between S2 and S3. Clusters DRG5 or H3 show an even more exotic behavior. Their unfolding curves show a dual nature: secondary peaks and unfolding humps (Fig. S14).

## Discussion

Given the high diversity of membrane proteins in cells, the unequivocal identification of any unlabeled protein from single cells is a daunting task. However, the present manuscript shows that by combing SMFS on native membrane patches with information present in databases, a probabilistic identification of groups of proteins is possible. Our approach provides usually less than 4 candidates (Fig. 5) for most of the identified clusters of F-d traces (p > 95%) and we were able to prove the validity of the identification by properly engineered constructs (Fig. 6).

The proposed pipeline offers also the possibility to enrich the structural information derived from Cryo-EM, in their native environment. SMFS cannot distinguish between a β-sheet and an α-helix, but it can determine with a good accuracy the unfolded domains(Engel and Gaub, 2008) of a membrane protein at room temperature (see Fig. 7). Cryo-EM cannot capture the structural heterogeneity of the poorly folded regions, while SMFS can provide important complementary information, for instance in loose connecting loops.

The pipeline, in addition, offers the possibility to obtain the unfolding profile of the membrane proteins from a limited amount of native material (membranes isolated from 1 to 10 cells). We have shown that the unfolding profile of membrane proteins can be used as a fingerprint to characterize the sample under investigation (see Fig. 2–3 and Fig. S12). We foresee that this approach can be extended to characterize, and eventually distinguish, membranes in cells in healthy and sick conditions where other methodology cannot be applied because of sample scarcity.

A major limitation of the pipeline — in its present form — is the possibility to merge in the same cluster the unfolding of proteins with a different molecular identity: indeed, from the Mass Spec data it is clear that different proteins can have the same — or approximately the same — molecular weight, a similar unfolding pattern and a similar total unfolded length Lc. This issue is rather significant for short proteins, i.e., those with values of Lc between 50 nm and 150 nm. In order to overcome this limitation, it will be desirable to couple SMFS with a device (e.g., solid state nanopores(Lee et al., 2018)) able to identify, even qualitatively, specific amino acids during the unfolding process. Another limitation is that a protein, in order to be identified with our approach, must be abundant: the number of distinct membrane proteins that are present, for example in hippocampal neurons is believed to be of order hundreds but with only few of them representing the majority of the expressed proteins. With our approach indeed we identify only 10 clusters.

This work ss a first step for the use of SMFS in native samples, but despite the limitations here reported, we were still able to classify tens of unfolding pathways, identify with high accuracy part of them, and we could also reconcile some contrasting cryo-EM structures for the molecularly identified proteins. We therefore believe that the proposed methodology represents a genuine step in the direction of single-cell membrane proteomics.

## Methods

All experimental procedures were in accordance with the guidelines of the Italian Animal Welfare Act, and their use was approved by the SISSA Ethics Committee board and the National Ministry of Health (Permit Number: 630-III/14) in accordance with the European Union guidelines for animal care (d.1.116/92; 86/609/C.E.).

### Cell preparation and culture

#### NG108-15

Mouse neuroblastoma NG108-15 cells were obtained from Sigma-Aldrich. The cells were grown in Dulbecco’s Modified Eagle Medium (DMEM, ThermoFisher) plus 10% Fetal bovine serum (FBS, Gibco), 100 U/ml Penicillin and 100 U/ml Streptomycin. The cells were cultured into a humidified incubator (5% CO2, 37 °C).

#### Hippocampal and DRG neurons

Hippocampal and DRG neurons were obtained from Wistar rats (P2-P3) as described in ref. (Galvanetto, 2018). In short, the animals were anesthetized with CO2 and sacrificed by decapitation. The dissociated cells were plated at a concentration of 4 × 10^4^ cells/ml onto glass round coverslips (170 μm in thickness) coated with 0.5 mg/ml poly-D-lysine (Sigma-Aldrich, St. Louis, MO, USA) for 1 h at 37°C and washed 3 times in deionized water. It is fundamental to obtain an optimal adhesion of the cells to prevent detachment in the next step (isolation of the cell membrane). The medium used for hippocampal neurons is in Minimum Essential Medium (MEM) with GlutaMAX supplemented with 10% Fetal Bovine Serum (FBS, all from Invitrogen, Life Technologies, Gaithersburg, MD, USA), 0.6% D-glucose, 15 mM Hepes, 0.1 mg/ml apo-transferrin, 30 μg/ml insulin, 0.1 μg/ml D-biotin, 1 μM vitamin B12 (all from Sigma-Aldrich), and 2.5 μg/ml gentamycin (Invitrogen). The medium used for DRG neurons is Neurobasal medium (Gibco, Invitrogen, Milan, Italy) supplemented with 10% Fetal Bovine Serum (FBS, from Invitrogen, Life Technologies, Gaithersburg, MD, USA).

#### Rods

Rod cells were obtained from adult male Xenopus laevis as described in ref. (Mazzolini et al., 2015). Under infrared illumination, the eyes of dark-adapted frogs after anesthesia with MS-222 were surgically extracted. Eyes were preserved in the Ringer solution (110 NaCl, 2.5 KCl, 1 CaCl2, 1.6 MgCl2, 3 Hepes-NaOH, 0.01 EDTA, and 10 glucose in mM; pH 7.8 buffered with NaOH), and hemisected under a dissecting microscope. The extracted retina was maintained in the Ringer solution.

### Cell transfection

NG108-15 cells were transiently transfected with 300 ng of each cDNA expression plasmids (see Supplementary Table 1) by using Lipofectamine 2000 Transfection Reagent (ThermoFisher) according to its handbook. Briefly, the plasmids (Supplementary Table 1) and the Lipo2000 were diluted into Opti-MEM Reduced Serum Medium (Gibco), respectively. 5 mins later, we added the diluted DNA to the diluted Lipo2000 to make the plasmid DNA-lipid complexes. After incubating 30 min, we plated the cells on the 12 mm round coverslips coated with 1x Poly-D-Ornithine (Sigma-Aldrich) in 12 well plate, and in the meanwhile, we added DNA-lipid complexes to the cells. We performed membrane isolation and confocal experiments about 56 hours after transfection.

### Isolation of cell membranes

#### Single-cell unroofing (for cell types that grow in adhesion)

The apical membrane of Hippocampal neurons, DRGs and NG108-15 cells were isolated with an optimized version of the unroofing method(Galvanetto, 2018). Briefly, additional empty glass coverslips (24 mm in diameter, 170 μm in thickness) were plasma cleaned for 15 seconds and broken in 4 quarters (with the use of the hands) in order to obtain optically sharp edges, as described in (Galvanetto, 2018). The coverslip quarters were immersed into 0.5 mg/ml poly-D-lysine for 30 minutes, and then they were immersed in deionized water for 10 seconds before use. A petri dish was filled with Ringer solution (3.5 ml) without any glucose, where the glass quarter was placed tilted of 7-15 degrees in the middle of it, supported by a 10 x 10 x 1 mm glass slice and Vaseline. The cover of the petri dish was then fixed on the stage of the AFM-inverted microscope setup (JPK Nanowizard 3 on an Olympus IX71).

The cell culture was then mounted on a 3D printed coverslip holder connected to the head stage of the AFM. The AFM head was put on top of the stage in measurement position. The cell culture was immersed into the solution and a target cell was identified and aligned with the underlying corner of the glass quarter. The cell culture was moved towards the corner of the underlying glass with the motors of the AFM until the target cell was squeezed and it doubled its area. At this point the cell is kept squeezed for 3 minutes, then a loaded spring under the AFM is released to abruptly separate the corner from the cell culture, and break the target cell membrane. The glass quarter with the isolated cell membrane was laid down and fixed on the petri dish. The medium was replaced with Ringer’s solution without exposing the cell membrane to the air.

#### Membrane isolation of non-adherent cells

Cells that do not grow in adhesion usually do not establish a tight binding with the substrate on top of which they are deposited. For these cells (e.g. rod cells), instead of unroofing, it is more reliable to break the cells with a lateral flux of medium(Clarke et al., 1975).

Isolated and intact rods were obtained by mechanical dissociation of the Xenopus retina in an absorption buffer (150 mM KCl, 25 mM MgCl2, and 10 mM Trizma base; pH 7.5); they were then deposited on cleaved mica as described in ref.(Maity et al., 2017). Incubated rods were maintained for 30–45 minutes over the mica in order to be adsorbed by its negatively charged surface. In the meanwhile, the position of the rods in the field of view of the microscope was annotated. The absorption buffer was substituted by a solution containing (in mM): 150 KCl, 10 Tris-HCl, (pH 7.5) and then a lateral flux of medium was applied to the rods until all the cell bodies were removed.

Experiments on the (almost) protein-free membrane of Oocytes, overexpressing CNG-N2b were made following the protocol of ref. (Maity et al., 2015).

#### Isolation of rod discs

Purification techniques with multiple centrifugations are usually required to isolate membrane-only organelles like rod discs or outer membrane vesicles(Thoma et al., 2018). Rod discs were obtained starting from the extracted retina as described in ref. (Maity et al., 2017). Briefly, discs were separated with two series of centrifugations of the sample overlaid on a 15-40% continuous gradient of OptiPrep (Nycomed, Oslo, Norway). 40 μl of the sample were diluted with 40 μl of absorption buffer, and incubated on freshly cleaved mica for 40 minutes. After 40 minutes, the incubation medium was removed and substituted with the solution used in the AFM experiments (150mM KCl, 10mM Tris-HCl, pH 7.5).

### Reconstitution of purified membrane proteins

Sthk channel was purified as described in ref.(Marchesi et al., 2018). bR and ChR1 proteins were bought from Cube-Biotech. The purified protein or they different mixtures (as indicated in the Results section) were brought at a total concentration of 0.5 mg/ml in a buffer containing 20mM HEPES, pH 7.8, 150mM KCl and 0.1% N-dodecyl β–D maltoside (DDM) and aliquoted into 100μL samples. A lipid mixture of 1,2-dioleolyl-*sn*-glycero-3-phosphocholine (DOPC), 1,2-dioleolyl-*sn*-glycero-3-phosphoethanolamine (DOPE), 1,2-dioleolyl-*sn*-glycero-3-phospho-L-serine (DOPS) at 8:1:1 ratio (Avanti Polar Lipids) was added at a lipid-to-protein ratio of 1 (wt/wt). The different ternary mixtures of protein-lipid-detergent were sonicated in an ice-bath sonicator for 2 min and subsequently equilibrated on an orbital shaker (200rmp) for 2h. Detergent was removed by hydrophobic adsorption adding twice ~5 mg of wet bio-beads SM-2 and equilibrating under gentle shaking (200rmp) for 1h at RT and overnight at 4°C, respectively. Before use samples were diluted 1:3 in adsorption buffer (20mM HEPES, pH 7.8, 150mM KCl)

The reconstituted membrane proteins were adsorbed on freshly cleaved mica for 30 min in a humid chamber. The samples were gently rinsed with imaging buffer (150Mm KCl, 10mM HEPES, Ph=7.8) and subsequently used for AFM imaging and unfolding.

### Gene silencing experiment

The genes of TRPC1/5/6 and TMEM16F were edited using CRISPR/Cas9 and their sgRNAs were cloned into LentiCRISPR-V2 puro vector (Supplementary Table 2). 3000ng of the indicated plasmids were transfected into the NG108-15 cells by using the lipo2000. 52h later after transfection, the cells were used for western blot. In order to select the silencing cell for single-cell unroofing, we treated the cells with culture medium containing 2 μl/ml puromycin(Jiao et al., 2018) after 22h transfection. After about 68h selection of puromycin, the alive cells were subjected to unroofing and the unroofed membrane were used for unfolding experiment.

### AFM imaging and Single-Molecule Force Spectroscopy (SMFS)

AFM experiments was performed using an automated AFM (JPK Nanowizard 3) with 50 μm long cantilevers (AppNano HYDRA2R-NGG, nominal spring constant = 0.84 N/m). We calibrated the AFM cantilevers in the experimental medium before each experiment using the equipartition theorem(Butt et al., 1995). The AFM experiments of Hippocampal neurons and DRGs were performed with Ringer’s solution (NaCl 145 mM, KCl 3 mM, CaCl2 1.5 mM, MgCl2 1 mM, HEPES 10 mM, pH 7.4); The AFM experiments of NG108-15 cell membranes were performed in the Ca2+-free Ringer solution supplemented with 10mM TCEP (Cat# C4706, Sigma). The TCEP can breakdown the disulfide bonds; Rod membrane and discs experiments were performed with 150mM KCl, 10mM Tris-HCl, pH 7.5. The fishing of the reconstituted membrane proteins (ChR1 and SthK) and bR was performed with imaging buffer (150mM KCl, 10mM HEPES, pH=7.8). All experiments were performed at 24 Celsius.

#### AFM imaging

The position of the cells before unroofing was annotated in the monitor of the computer in order to start the AFM imaging where the cells was in contact with the substrate (cell membrane is not visible in bright-field). The membrane obtained with single-cell unroofing (hippocampal neurons, DRG and NG108-15) can easily be found in proximity of the glass corner (~80% success rate). In the case of the rod membrane (non-adherent cells), usually different positions need to be scanned before finding a patch of membrane. Rod discs, and the reconstituted protein patches can be identified only via AFM imaging. We performed imaging both in contact mode (setpoint ~0.4 nN) and intermittent-contact mode (lowest possible), but the intermittent-contact mode if preferable because it does not damage the border of the patches of membrane.

#### AFM-based SMFS (protein unfolding)

we performed automated SMFS on top of the imaged membranes by setting grid positions for the approaching and retraction cycles of the cantilever. All experiments were performed with a retraction speed of 500 nm/s (for hippocampal neurons, DRG, Rods and Discs), 600nm/s (for NG108-15 cells), 1000nm/s (for bR, ChR1 and SthK). The membrane proteins present in the sample were attached non-specifically to the cantilever tip by applying a constant force of ~1 nN for 1 second between the AFM tip and the cytoplasmic side of the membrane. This method proved to work with different membrane proteins(Tanuj Sapra et al., 2006; Thoma et al., 2017, 2012), and to allow a higher throughput compared to methods that involve a specific attachment between the tip and the protein(Cisneros et al., 2005; Kedrov et al., 2004; Müller et al., 2002; Oesterhelt et al., 2000; Sumbul et al., 2018a, 2018b). Also, in order to demonstrate the sawtooth patterns truly represent the unfolding of membrane proteins, we performed unfolding experiment on the top of the coverslip coating with the poly-D-lysine.

### Western blot

The transfected NG108-15 cells were harvested in the lysis buffer(150 mM NaCl, 1.0% Triton X-100, 0.1% SDS, 0.5% sodium deoxycholate, 10 mM NaF, 1 mM Na3VO4, 1.0% NP-40, and 40mM Tris-HCl, pH=8.0) supplemented with protease inhibitor(Cat# 469311001, Roche) on ice. The samples were boiled at 100 °C for 30 mins and then the supernatants were loaded in 10% SDS PAGE gel for gel electrophoresis. The proteins on the gels were transferred onto the PVDF membranes (Cat# 88520, ThermoFisher) and these membranes were blocked with 5% nonfat dry milk. The membranes were incubated with the corresponding primary antibodies 1:5000 for α-Tubulin (Cat# T8203, sigma); 1:500 for TRPC1 (Cat# ACC-101, Alomone Labs)/5 (Cat# ACC-020, Alomone Labs)/6 (Cat#ACC-120, Alomone Labs) and 1:200 for TMEM16F (Cat# ACL-016, Alomone Labs) at 4°C overnight with gentle shaking. Secondary antibodies Goat-anti-rabbit HRP (Cat# P0448, Dako) and Goat-anti-mouse (Cat# P0447, Dako) 1:5,000 in 5% BSA were used. Membranes were washed with PBST three times and developed with an ECL-HRP (Cat# WBULS0100, Millipore) system. The gray value ratios of protein bands were quantified using ImageJ software.

### Confocal experiments

#### NG108-15 whole cells

The transfected cells and non-transfected cells were fixed by using 4% PFA for 60 min at RT. The transfected cells were stained with Hoechst (Life Technologies). The non-transfected cells were permeated by 0.05% TritonX-100 for 4 min and blocked by 10% FBS+5% BSA for 90 min, following by the incubation of corresponding primary antibodies (1: 250) at 4°C overnight. The samples were rinsed with pre-cold 1xPBS 3 times and incubated with the secondary antibody Alexa 594-labeled goat anti-rabbit (1:800, Cat# A11037, Invitrogen) at RT for 90 min. After staining the samples with Hoechst, the samples were tested by Nikon A1R microscope with 60× oil immersion objective (NA 1.40). The results were analyzed with the ImageJ software.

#### NG108-15 cell membranes

The NG108-15 cells were stained with 200 μM SiR-Actin, 200 μM SiR-Tubulin (Spirochrome) or 25 nM MitoTracker® Red FM (ThermoFisher) for 30 min in a humidified incubator (5% CO2, 37 °C). The stained or transfected cells were unroofted according to the method as mentioned above. Then the NG108-15 cell membranes were tested by Nikon A1R microscope with 20× objective, followed by the imaging scan by AFM. The results were analyzed with the ImageJ software.

### Molecular visualization

We aligned the current protein structures of TRPC superfamily using the PyMOL software, such as human TRPC3, mouse TRPC4, mouse TRPC5 and human TRPC6. Their codes in PDB are 5zbg, 5z96, 6aei and 5yx9, respectively. We zoomed the C-terminal part (from S5 to the C-termini) of these structures and find more details.

## Acknowledgments

We thank Alexander Bruce for useful comments on the manuscript. We thank dr. Kosaku Shinoda for support in the emPAI calculation. We thank Prof. Anna Menini who provided the mTMEM16A-GFP and mTMEM16F-GFP plasmids. We thank Prof. Guidalberto Manfioletti who provided the peGFP-N1 plasmid.

## Supplementary Information

- Supplementary Methods
- Figs. S1 to S14
- Tables S1 to S2

### Automatic classification of SMFS data

The selection of the F-D curves that represent the unfolding of membrane proteins is usually based on the search for a specific pattern of unfolding in the SMFS data, after a filtering based on the length of the protein under investigation^1,2^. In the case of a native preparation (like ours) that contains a mixture of unknown proteins a) the filtering based on the distance cannot be applied and b) the number of specific patterns to be found is unknown. In order to find recurrent patterns of unfolding in a SMFS dataset we developed an algorithm that consists of five major blocks (Fig. S11 a). In the first block, the parts of the F-D curves not related to the unfolding process are removed, and a coarse filtering aimed at the detection of spurious traces is performed. In the second block, a quality score based on the consistency of the experimental data with the worm-like chain (WLC) model is computed and assigned to each trace. This score is used to select physically meaningful traces for further analysis. In the third block, distances between pairs of traces are computed to assess their similarity. The distances are used in the fourth block for density peak clustering. The fifth and final block consists in the refinement and possibly in the merging of some of these clusters. In what follows we provide a detailed overview of each block.

#### Block 1: filtering

The standard F-D curve preprocessing was applied to all the data within ‘Fodis’^3^. The zero of the force of the curve was determined averaging the non-contact part (baseline after the final peak) and subtracted to all the points of the curve. The piezo position was transformed in tip-sample-separation considering the contribution of the bending of the tip to the extension of the polymer. Given that the F-D curves are subject to noise (due to thermal fluctuations, coming from the instrument, etc.), we smooth the original signals through interpolation on a grid with width δ_interp_= 1 nm.

A curve is discarded if it does not contain a:

- detectable contact point (i.e. a transition from negative forces to positive forces in respect to the baseline set at zero force);
- if the points occupy force ranges over 5000 pN;

Some of the F-D curves show deviations from the horizontal zero-force line in the non-contact part (wavy final part due to imperfect detachment of the polymer or other noise from the environment). We detect and discard these traces by computing the standard deviation of the tails from the zero-force line. If it exceeds two times σ_NOISE_ (average standard deviation of the baseline of the batch of curves) the trace is discarded.

#### Block 2: Quality score

The quality score is used to refine selection of traces with high information content vs. noisy traces. It is based on the description provided by the worm-like chain (WLC) model, which is the standard model in the analysis of SMFS data^4^. The WLC model implies the equation:

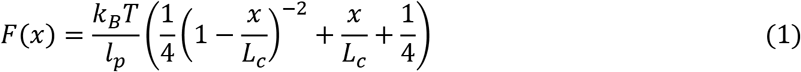

where *F* is force, *x* is extension, *k_B_* is Boltzmann’s constant, *T* is temperature, *l_p_* is persistence length and *L_c_* is contour length. Each unfolding curve in the trace is fitted with the WLC equation and a *L_c_* value, corresponding to the length of the unfolded protein domain is obtained. The *L_c_* values are computed by solving equation (1) for each *x* and *F*. An appropriate value for the persistence length *l_p_* for membrane proteins is 0.4 nm as reported in ref. ^4^. The WLC model is applicable in the force range 30-500 pN^5^. Once we compute the *L_c_* values, we can build a *L_c_* histogram. Normally, the *L_c_* histogram describing a successful unfolding experiment is characterized by the presence of a few maxima separated by deep minima. We implement these features in the definition of our quality score to distinguish meaningful F-D curves.

An important parameter is the bin width of the *L_c_* histogram. If the bin width is too small the histogram is noisy; if the bin width is too large, peaks corresponding to the unfolding of different domains might be merged. We use bin width 8 nm which is an efficient value for evaluating the goodness of a curve and it allows to consider also curves that deviate from the WLC model (*l_p_* =0.4 nm) but that contain information. Furthermore, the choice of such large bin width is based on visual inspection of the histograms of proteins with known structure. Once the *L_c_* histogram is built, we detect all maxima and minima. A maximum is meaningful if it is generated by more than 5 points and it includes more than 1 % of the force measures of a trace.

For each maximum in the *L_c_* histogram, we compute a score *W* quantifying the consistency of the peak with the WLC model. A high-quality peak is clearly separated from other peaks of the histogram, therefore it should be surrounded by two minima. We define 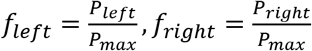 where *P_max_, P_left_* and *P_right_* are the probability densities of the maximum, of the left and the right minima. Ideally, 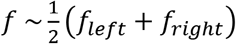 should go to 0. We define the peak score as *W* = *exp*(−2*f*^2^). According to this definition, if *P_left_* = 1, *P*_right_ = 2 and *P_max_* =16, W=0.98. Whilst if *P_left_* = 13, *P_right_* = 14, the peak doesn’t fit well with the WLC model and W=0.24.

Once a score is computed for each relevant peak in the *L_c_* histogram, that score is assigned to all points in the corresponding trace. This is accomplished in two steps: first, the peak score is assigned to all points in the histogram belonging to that peak. Second, to all points with force values below 30 pN, for which an *L_c_* values cannot be computed due to the model’s limitations. To these points, we assign the score of the first successive point with force larger than 30 pN. This criterion applies only to points within 75 nm from the last point assigned to the peak. The peak width value is selected by visual inspection of traces, evaluating the maximum width of their force peaks.

The quality score of a trace, *S_w_*, is the sum of the scores for all points in the trace. The higher the global score, the higher the trace quality. We use the ratio between the quality score and the trace length to select high quality traces. If this ratio is below 0.5, we discard the trace. We assume that if more than half of the trace is inconsistent with the WLC model, it is a low-quality trace and as such we exclude it from the analysis. While if more than half of the trace is in good agreement with the WLC model, it is possibly a meaningful trace.

We point out that the goal of blocks 1–4 is only to find dense recurrent patterns in the SMFS data: in block 5 we reevaluate the F-D curves to form the selections shown in Fig. 3 of the main text.

#### Block 3: Computing distances

In block 3 we quantify the similarity between the traces in order to find the recurrent pattern of unfolding within the data. To accomplish this goal, we use a modified version of the distance introduced by Marsico et al.^1^. This distance is defined using the dynamic programming alignment score computed for a pair of traces. For two traces, *a* and *b*, the distance *d_ab_* is simply:

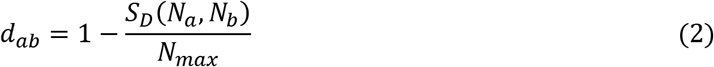

where *S_D_* (*N_a_, N_b_*) is the global alignment score, *N_a_* is the length of trace *a, N_b_* is the length of trace *b*, and *N_max_* is the maximum length between the two. We have modified the match/mismatch scoring function used by Marsico et al as follows:

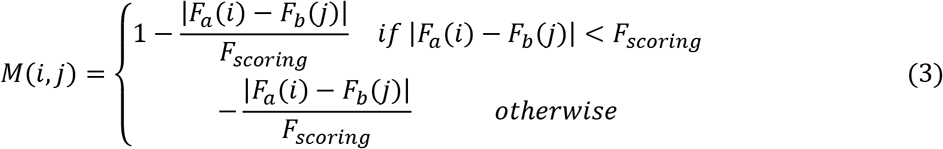

where *F_a_*(*i*) and *F_b_*(*j*) are the forces in points *i* and *j* in traces *a* and *b*, and *F_scoring_* = 4*σ_NOISE_*. In the work done by Marsico et al, *F_scoring_* is replaced by Δ*F_max_*, which is the average of the maximum force values in the two traces. When two widely different traces have high Δ*F_max_* their distance will be lower with respect to two traces with low Δ*F_max_* but overall higher level of similarity. Namely, the distance magnitude depends on the Δ*F_max_* value and traces with high Δ*F_max_* have by definition lower distance values. It is important to note that this problem did not occur in Marsico’s work since the Δ*F_max_* values were uniformly distributed for all traces.

In order to gain computational efficiency, the distance is computed only for traces which differ by no more than 2 peaks in the *L*_c_ histograms or by no more than 20 % in their trace length difference.

#### Block 4: Density peak clustering

The density peak clustering (DPC) algorithm^6^ is used for clustering. This choice is appropriate given that a fraction of traces in the analyzed datasets correspond to statistically isolated events and DPC automatically excludes the outliers. DPC can be summarized in the following steps:

1. We compute the density of data points in the neighborhood of each point using the *k*-nearest neighbor (*k*-NN) density estimator^7^. The density is the ratio between *k* and the volume occupied by the *k* nearest neighbors:

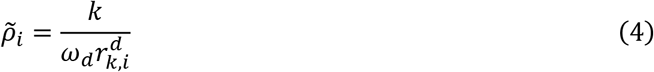

where *d* is the intrinsic dimension (ID) of the dataset^8^, *ω_d_* is the volume of the *d*-sphere with unitary radius and *r_k,i_* is the distance of point *i* from its *k*-th nearest neighbor. In DPC it is the density rank which is relevant for the final cluster assignation. Therefore, without loss of generality, we compute the density using the following equation:

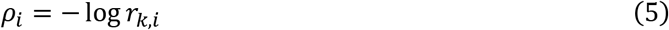

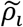 and *ρ_i_* are related by a simple monotonic transformation and thus, have the same rank. By using equation (4) we don’t have to compute the intrinsic dimension of the dataset. In order to assign bigger weight to high quality traces, we multiply *ρ_i_* by the score-length ratio of trace *i*.
2. Next, we find the minimum distance between point *i* and any other point with higher density, denoted as *δ_i_*:

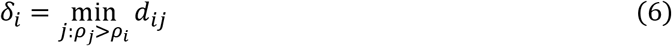

where *d_ij_* is the distance between points *i* and *j*. *δ_i_* is used to identify the local density maxima.
3. We identify the cluster centers as density peaks, e.g. points with high values of both *ρ_i_* and *δ_i_*. For each point we compute the quantity *γ_i_* = *ρ_i_δ_i_*. Points with high values of *γ_i_* are good cluster center candidates. We sort all points by the value of *γ_i_* in descending order. The first point is a cluster center. The second point is a cluster center unless its distance from the first point is smaller than *r_cut_* = 0.3 (which represents the distance below which on average two traces are considered as the same pattern). Regarding the third point, it is a cluster center if it is at a distance smaller than *r_cut_* from the preceding two points. Following the same logic, all the points are assessed and all cluster centers are identified.
4. All points that are not cluster centers are assigned to the same cluster of the nearest point with higher density.

#### Block 5: Refinement and merging

The previous blocks, from 1 to 4, were optimized for finding the centers of dense patterns of unfolding in the SMFS data, but not for finding the borders of the clusters. To solve this issue, i.e. finding the F-D curves that are similar to each pattern of unfolding, we used the conventional definition of similarity (degree of superposition of F-D curves in the Force/tip-sample-separation plane) automated in the Fodis software in the tool ‘fingerprint_roi’^3^.

In brief, we superimposed each cluster center with its two closest neighbors creating the effective ‘area of similarity’ (AoS) for each cluster. The AoS is defined as the area generated by all the points of the three curves above 30 pN and before the last peak (see Supplementary Fig. 11 b), each point forming a square of 5 nm x 5 pN. Then, the SMFS curves are preliminary filtered based on their length with their final peak falling between 0.7 x *L* and 1.3 x *L* (with *L* length of the cluster center). Each of the remaining F-D curves is compared with the AoS, and the number of its points that fall within the AoS is annotated: this number constitutes the similarity score. As depicted in Supplementary Fig. 11 c, the plot of the scores in descending order interestingly forms a line with two different slopes. The change of the slope empirically defines a threshold that reflects the limit of similarity for each cluster. If two clusters share more than 40% of the traces above the threshold, they are considered the same cluster, thus merged (all the merges are reported in Supplementary Fig. 11 d).

### Formal derivation of the Bayesian identification of the clusters

Bayesian inference is widely used in modern science^9,10^ because it allows to univocally determine the level of uncertainty of a hypothesis^11^. We used the same framework to determine the molecular identity of the unfolding clusters. In the most general terms, we observed the unfolding cluster *C_x_*, and we want to find the probability that the unfolding of a certain protein *Prot_A_* corresponds to the unfolding cluster *C_x_*, i.e. we want to find the posterior probability *P(Prot_A_*|*C_x_*). In the form of the Bayes theorem:

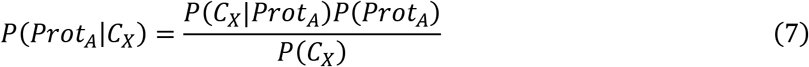

where *P*(*Prot_A_*) is the prior, i.e. the probability of *Prot_A_* to be in the sample; *P*(*C_x_*|*Prot_A_*) is the likelihood, i.e. the likelihood to find a cluster with the features of *C_x_* coming from the unfolding of *Prot_A_*; and *P*(*C_X_*) is the normalizing factor. In the case of a classical experiment with a single purified protein, *P*(*Prot_A_*|*C_x_*) is assumed to be equal to 1, but this is not the case for a native environment where there are *Prot_B_*, *Prot_C_*, etc.

The observables of an unfolding cluster for which we determined the likelihood functions are the contour length of the last detectable peak *LC_max,Cx_* (~ length of the F-D curve), the peak profile of the cluster in the Lc space *Peaks_Cx_* (also called unfolding barriers), and the average unfolding force of the detected peaks 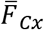, but the method is modular and therefore it will incorporate also other observables when available. The equation (7) becomes:

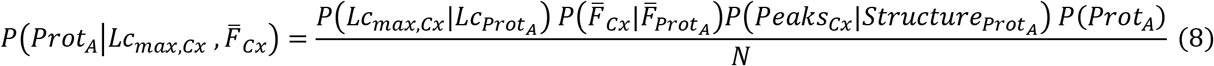

where 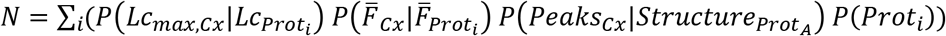 is the normalizing factor that takes into consideration all the proteins *Prof* present in the sample. In the next paragraphs we will describe the determination of the numerator of equation (8).

#### *Determination of prior P*(*Prot_A_*)

The most crucial part of the method is the determination of the list of proteins present in the sample, together with all their properties (length, abundance, secondary structure, topology, etc.). To do so we combined the Mass Spectrometry results of the cells under investigation^12–14^ with other structural and topological information available in Uniprot and PDB. The crossing of the databases is done thanks to the unique Uniprot identifier. The complete list of proteins of Hippocampal neurons, Rod outer segments and Rod discs with the information necessary for the Bayesian inference are shown in the Supplementary Data of the article. In case the data of the species of interest are not available, cross species proteomic analysis demonstrated that the majority of proteins are conserved in terms of relative abundance^15,16^. *P*(*Prot_A_*) is the probability of finding *Prot_A_* and not *Prot_B_*, *Prot_C_*, etc., which corresponds to the normalized relative abundance of *Prot_A_* in the sample—a parameter that is usually calculated in mass spectrometry analysis. Indeed, *in silico* calculation of abundances gives rather trustworthy values:

1. the most accurate option is the emPAI^17^;
2. if the emPAI is not available, the second best option is the spectra counting for each peptide (PSM)^18^;
3. if the PSM is not available, the sequence coverage can be used as loose estimation^19^.

We used the emPAI for hippocampal neurons and rods; for the discs, the emPAI does not give accurate values because of the extreme concentration of Rhodopsin, therefore we used the abundances obtained with other quantitative methods^20^.

We demonstrated in Supplementary Fig 1, 2 and 3 that the isolated paches of membrane contain the membrane proteins of the original cells but not the cytoplasmic proteins, therefore we created an additional binary variable *ismembrane* for each protein, equal to 0 if the protein is not a membrane protein, 1 otherwise. This information is extracted from the annotation in the Uniprot database. The final prior is:

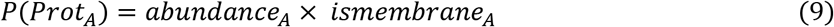

#### *Modeling the conditional probability P*(*Lc_max,Cx_*|*Lc_Prot_A__*)

The F-D curves encode a reliable structural information, that is the total length of the unfolded protein^21^. We revisited 14 published unfolding clusters of membrane proteins^21–33^ plus our own experiments, and that allowed us to create the conditional probability function for the observable *Lc_max,Cx_* as shown in Fig. 4 b–c. The distribution of the effective unfolding lengths forms a Gaussian bell centered at *0.89Lcp_rotA_* with a standard deviation of *σ_effective_* = 0.05*Lc_Prot_A__*. The final standard deviation is obtained combining *σ_effective_* with the error due to the unperfect determination of the persistence length *σ_p_* = 0.07*Lc_Prot_A__*, and the error of the *Lc_max,Cx_* that is *σ_Lc_* = 0.05*Lc_Prot_A__*, so we obtained 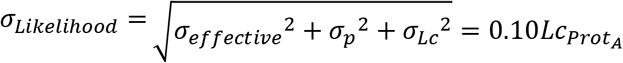. This prior is determined without considering any assumption on the pinning point of the protein (i.e. the position of the amino acid fixed at the level of the membrane that determine *Lc_max,Cx_*), just considering the experimental data.

Most of these experimental data comes from membrane proteins that do not have a final cytoplasmic/extracellular domain. These data show also that the 0.89*Lc_Prot_A__* correction of Fig. 4 c originates from a pinning point that can occur anywhere between the last amino acids of the last transmembrane domain and the penultimate loop.^25,30,31^ There are only few examples of SMFS of proteins with a large cytoplasmic domain at one terminal^33,34^, and these data suggest that the pinning point to the membrane or to another protein can be i) at the terminus end of the protein, ii) everywhere along the domain, or iii) at the end of the transmembrane segments. In other words, the pinning point of a protein that has a final domain cannot be firmly established if not within the domain range. Therefore, for the proteins that have topological annotations, we replaced the Gaussian prior described above with a non-informative prior normalized to *1* and constant between (*L_protein_* – *L_final domain_*) < *Lc_max,Cx_* < *L_protein_*, zero elsewhere.

We use this non-informative prior only for proteins with *L_final domain_* ≦ 2*σ_Likelihood_*; for proteins with small final domains *L_final domain_* < 2*σ_Likelihood_*, we used the Gaussian prior centered at 0.89*Lc_Prot_A__*. We chose the threshold of 2*σ_Likelihood_* in order to take into account the uncertainty on the position of the pinning points and the errors *σ_p_* and *σ_Lc_* that are still present even in proteins with small final domains.

#### *Determination of the conditional probability P*(*Peaks_Cx_*|*Structure_Prot_A__*)

Empirical evidence^21–33^ and simulations^35^ suggests that the occurrence of the peaks is more likely in presence of unstructured regions, i.e. in the intracellular or extracellular loops of the membrane proteins (see Fig. 4f). We used this information to evaluate the probability that the structure of the *Prot_A_* unfolds with the unfolding pattern of cluster *C_x_* (position of the peaks). To do so, we calculated the cross-correlation between *Peaks_Cx_* i.e. the global histogram of cluster *C_x_* (of point between 40 pN and 100 pN for normalization porpuses) and *Structure_Prot_A__* i.e. the profile of gaussian bells centered in the center of the loops of *Prot_A_* and with a FWHM of 15 nm which is an average estimate of the distance between two loops (see Fig. S13). If no structure is available we cross correlated a flat line. Both *Structure_Prot_A__* and *Peaks_Cx_* are normalized to have a total area under the profile equal to 1. Then we assigned to *P*(*Peaks_Cx_*|*Structure_Prot_A__*) the maximal value of the cross correlation with a relative lag of ±15 nm to allow some freedom for fine alignment.

#### Determination of the conditional probability 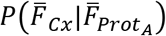

The force necessary to unfold a protein domain depends on the stability of the domain itself, α-helices and β-sheets are unfolded at different force levels as shown in Fig. 4 d. We revised the unfolding forces of 22 proteins^21–33,36^ and we used as 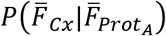 the smoothed trend line of the distribution (Fig. 4 e of the main text).

---

**Figure S1.**
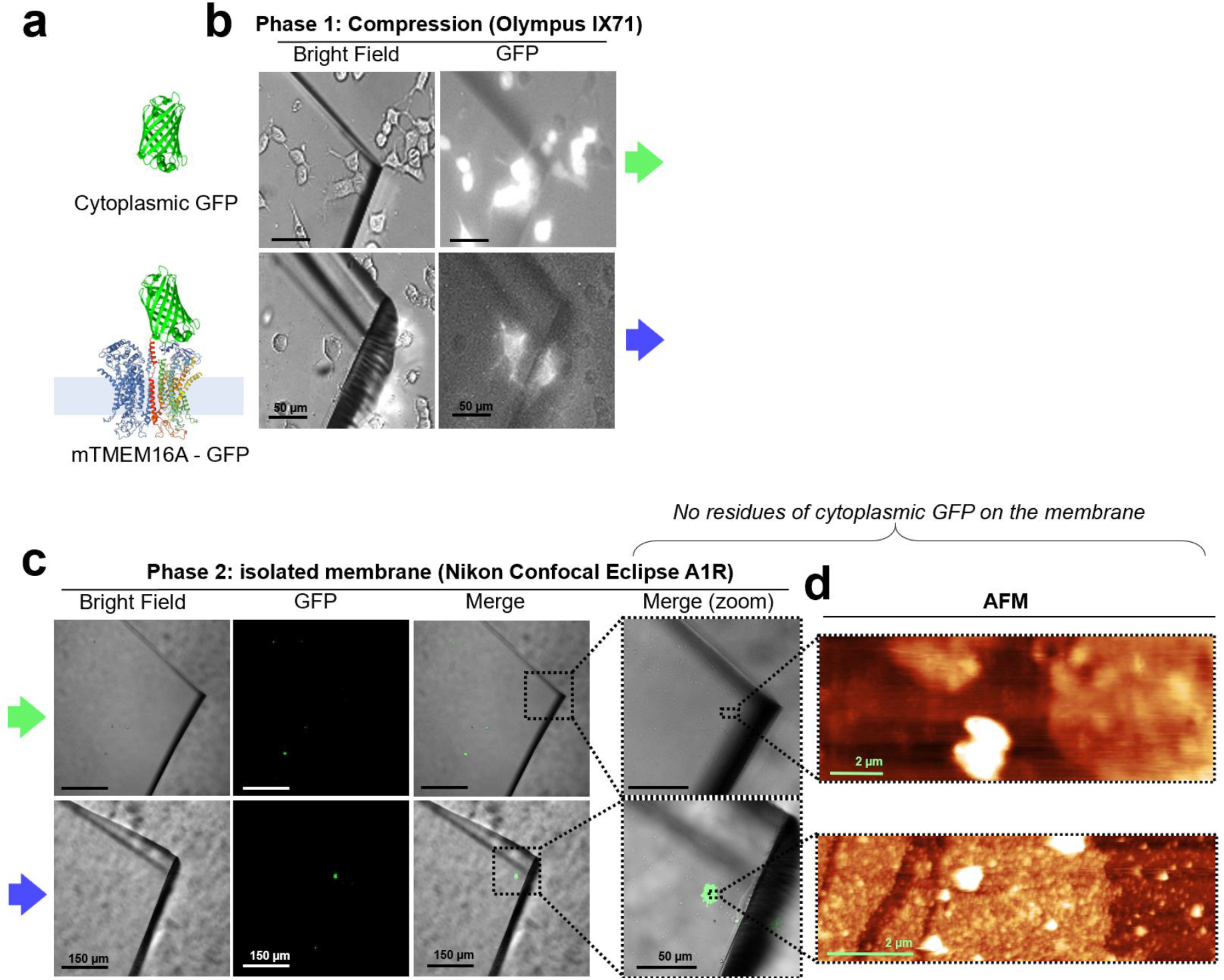
In NG-108 cells, membrane proteins remain into the membrane after unroofing, while cytoplasmic proteins do not. **a**, (top) cytoplasmic GFP and (bottom) mTMEM16A-GFP overexpressed in NG-108 cells. **b**, bright field and fluorescence images taken during the compression on the inverted microscope-AFM system. **c**, images taken with a confocal microscope of the coverslip quarter after unroofing; the rough surface under the glass is due to the Vaseline layer used to fix the coverslip quarter. No residues of cytoplasmic GFP are found in the top row. **d**, AFM images of the area in the right most panel of **c** showing the presence of membrane patches in both the samples.

**Figure S2.**
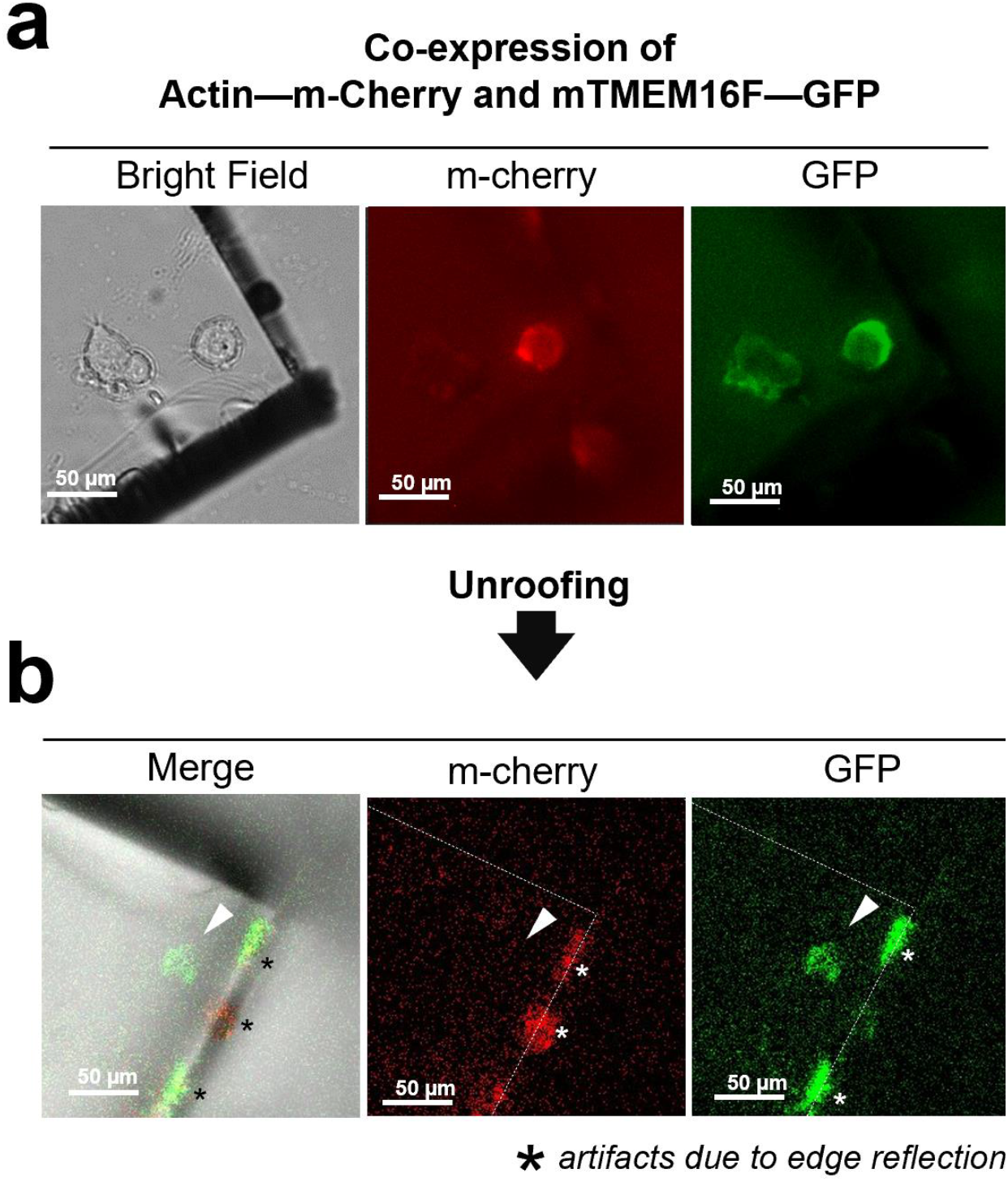
Absence of cytoplasmic proteins after unroofing of NG-108 cells. **a:** before unroofing. From left to right: bright field image showing a piece of glass over a neuroblastoma cells transfected with actin labelled with m-Cherry and the membrane protein TMEM16F tagged with GFP; in the center an image of the red fluorescence emitted by the labelled actin; in the right an image of the green fluorescence emitted by the TMEM16F-GFP. **b:** after unroofing only the green signal from TMEM16F is detected (right panel) and no red fluorescence is observed (center panel and the location indicated by the white arrowhead). * (asterisks) indicate the fluorescence due to edge reflections. The green signal is detected exactly where the neuroblastoma cell was located (left panel).

**Figure S3.**
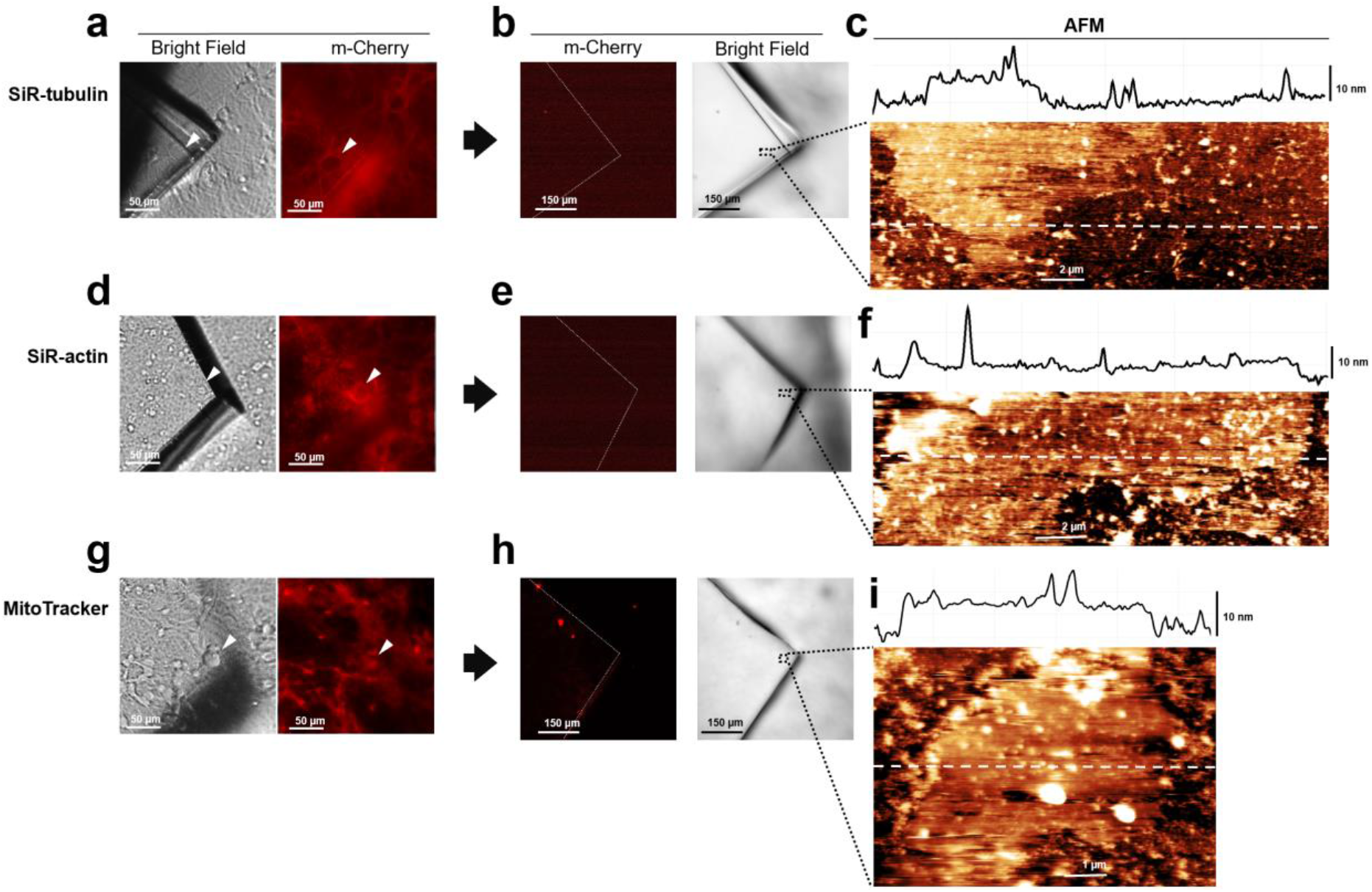
Absence of cytoplasmic proteins in unroofed membranes in hippocampal cells. **a:** before unroofing. Bright field image of the piece of glass over a neuroblastoma cell. Red fluorescence emitted by SiR-tubulin**. b:** after unroofing. No red fluorescent signal detected. **c**: AFM imaging of region indicated in **b** showing the presence of a piece of membrane. **d, e** and **f** as in **a, b** and **c** but for Sir-Actin. **g, h** and **i** as in **a, b** and **c** but for Mito Tracker labelling mitochondria.

**Figure S4.**
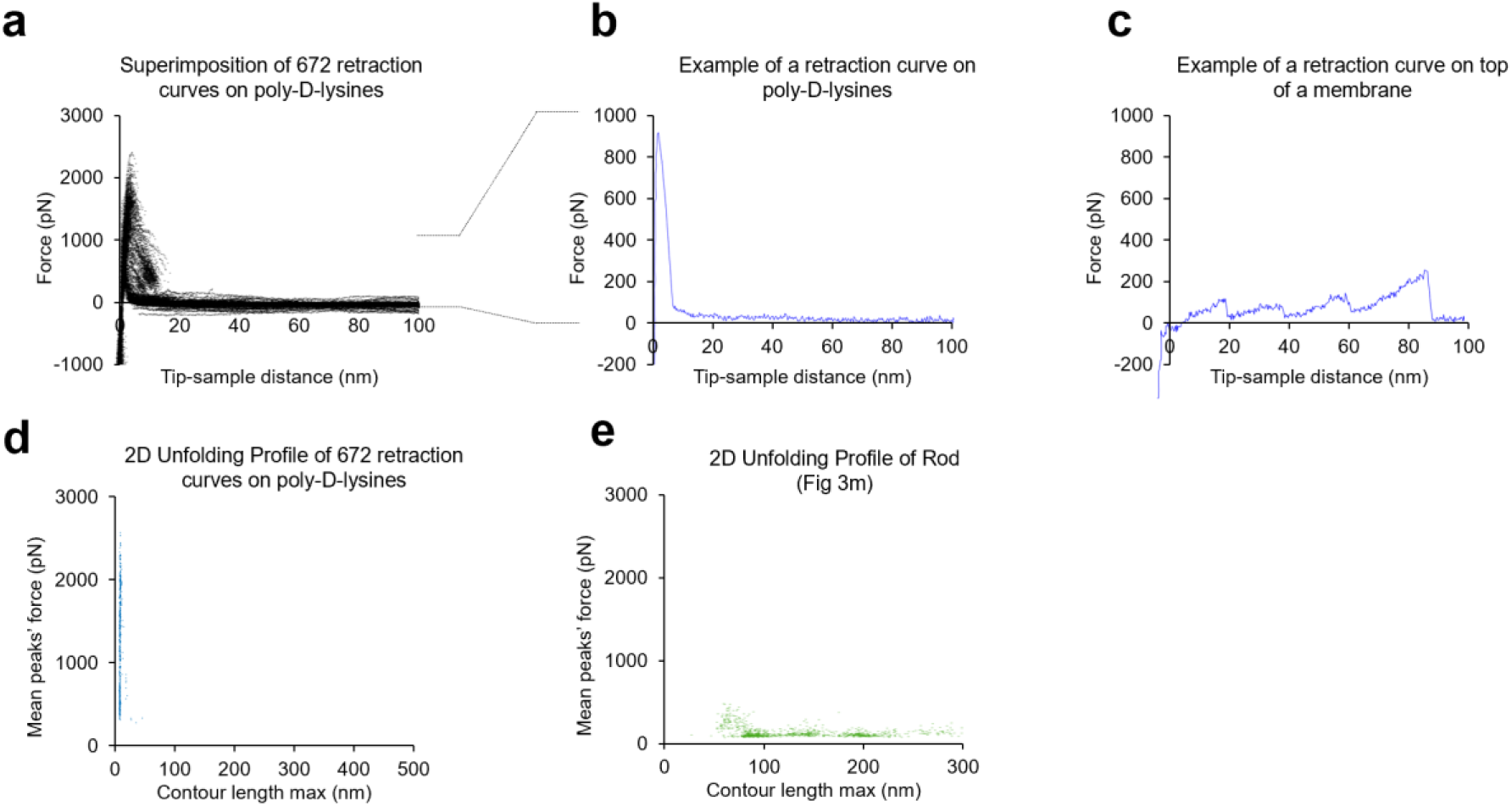
Absence of Poly-D-lysine contamination in the SMFS data from unroofed cell membranes. **a** 672 F-d curves obtained from a cover slip just coated with Poly-D-Lysine and in the absence of any cell. **b**: example of one F-d curve from **a**. **c**: one example of a F-d curve obtained in the presence of a biological membrane. **d**: plot of the mean force against maximum contour length of the traces obtained from the experiment shown in **a**; **e**: as in **d** but for F-d curves obtained from rod membranes.

**Figure S5.**
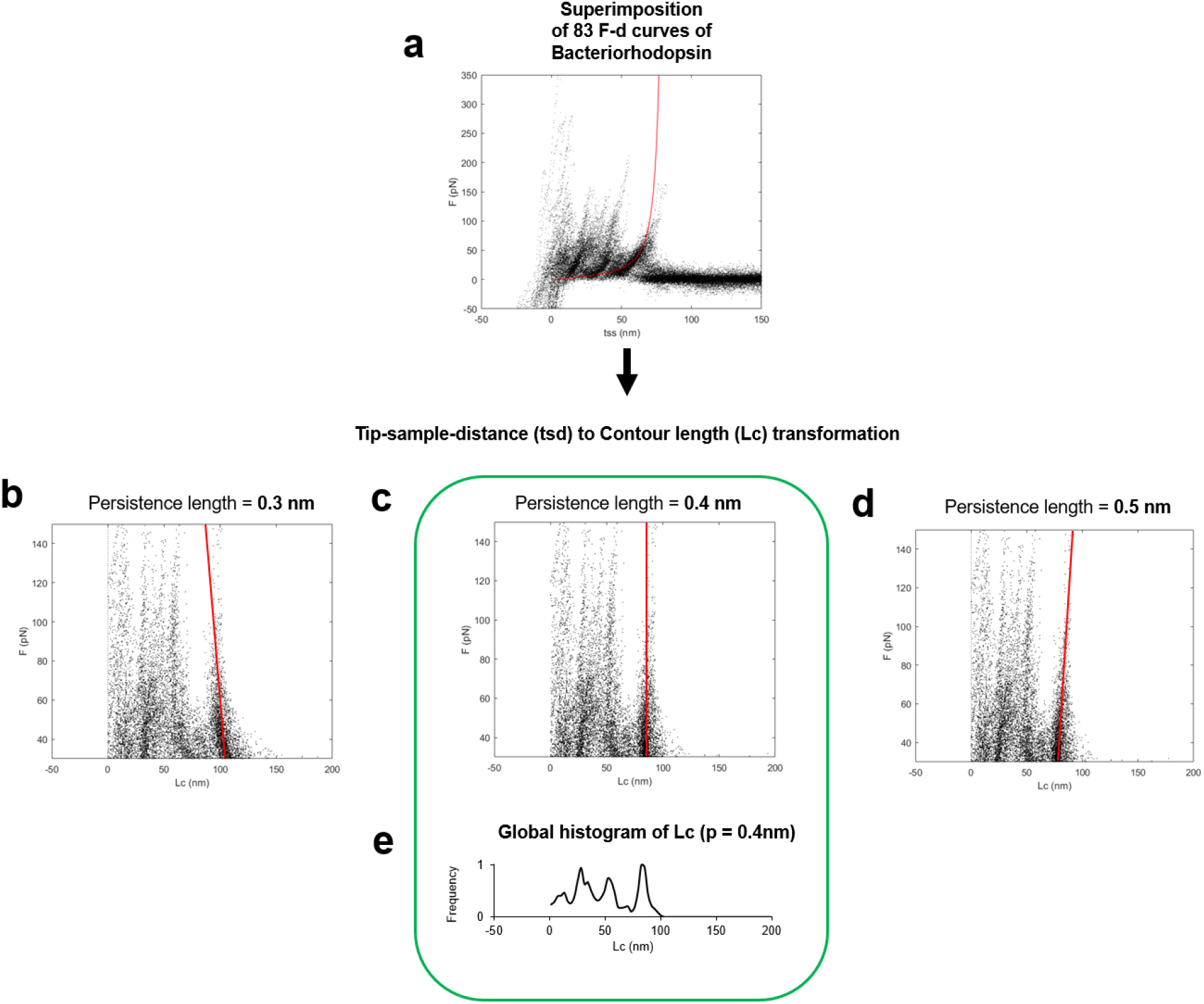
Determination of the optimal persistence length of the detachment peak used for the total contour length calculation (Lc max). **a:** superimposition of 83 F-d curves of Bacteriorhodopsin (the red line is the worm like chain curve with persistence length = 0.4 nm and contour length = 85 nm). **b, c, d:** Transformation of the ordinates of the points in **a** to the Contour length space (Lc) with persistence length of 0.3 nm in **b,** 0.4 in **c** and 0.5 in **d**. Solid red lines indicate the best fit of the points belonging to the last unfolding peak: 0.4 nm of persistence length is the correct/optimal value because in the contour length space, the force points representing the unfolding of the polymer with a constant contour length must lay on a vertical line. **d:** Global histogram of contour length of **c**.

**Figure S6.**
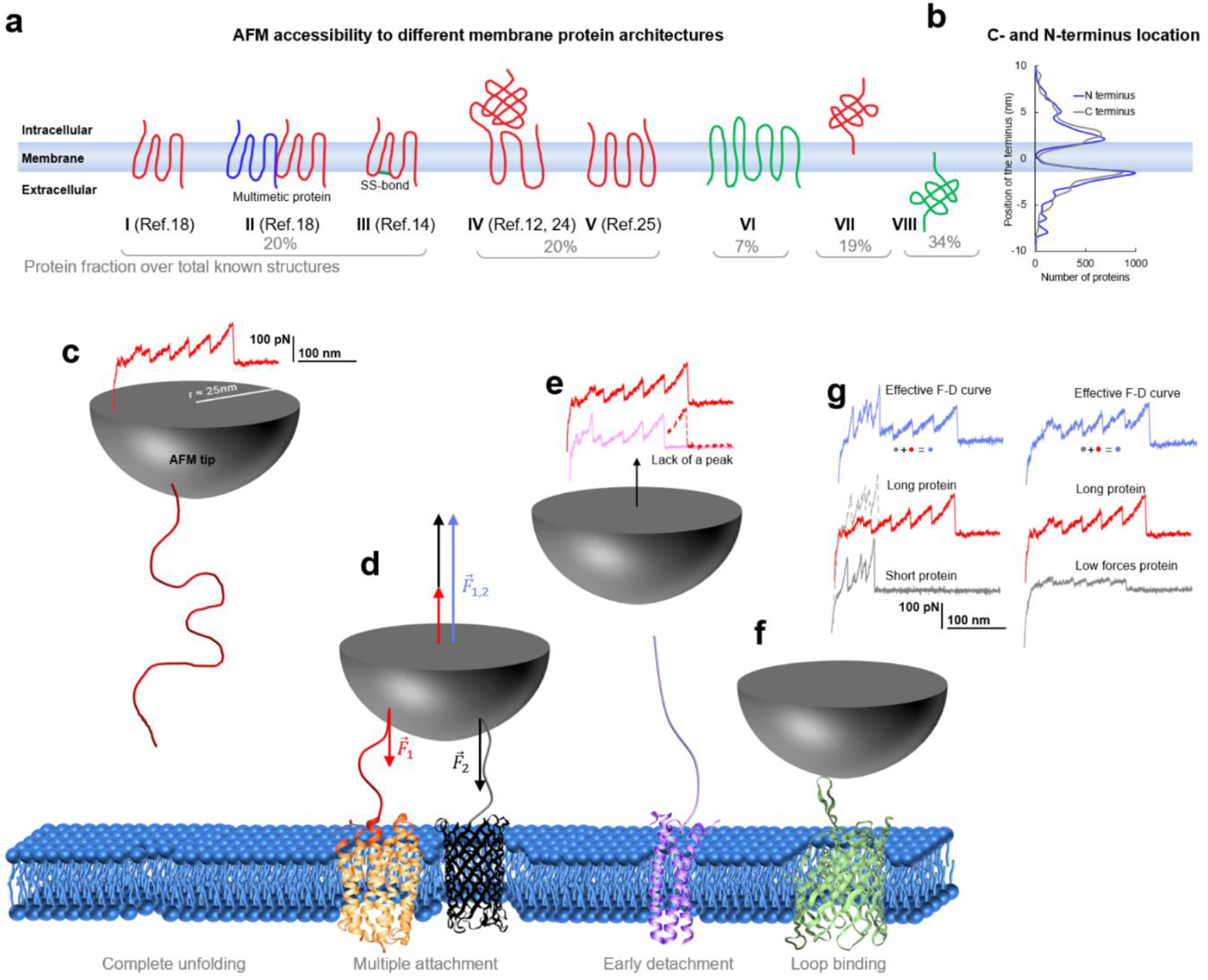
Membrane proteins architectures. **a**, eight classes of membrane proteins and their fraction obtained from data present in the PDB-OPM. **b**, distribution of the position of the C and N termini relative to the center of the cell membrane along the axis perpendicular to the membrane. **c**, Cartoon representing the complete unfolding of a membrane protein and its F-d curve, **d**, simultaneous unfolding of two proteins and the resulting involved force which is the arithmetic sum of the one necessary to unfold one protein ant the other. **e**, incomplete unfolding of a protein leads to a similar but shorter unfolding curve, **f**, unfolding from a cytoplasmic loop. **g**, idealized decomposition of F-D curves of a multiple unfolding/ unfolding from a loop. Effective F-d curves are affected by multiple binding: If one of the two protein is short, the unfolding of the longer one can still be recognized although it will differ in the intersection part. If one protein unfolds with lower forces that the other, still the patter of the second one can be recognized, but it will be noisy.

**Figure S7.**
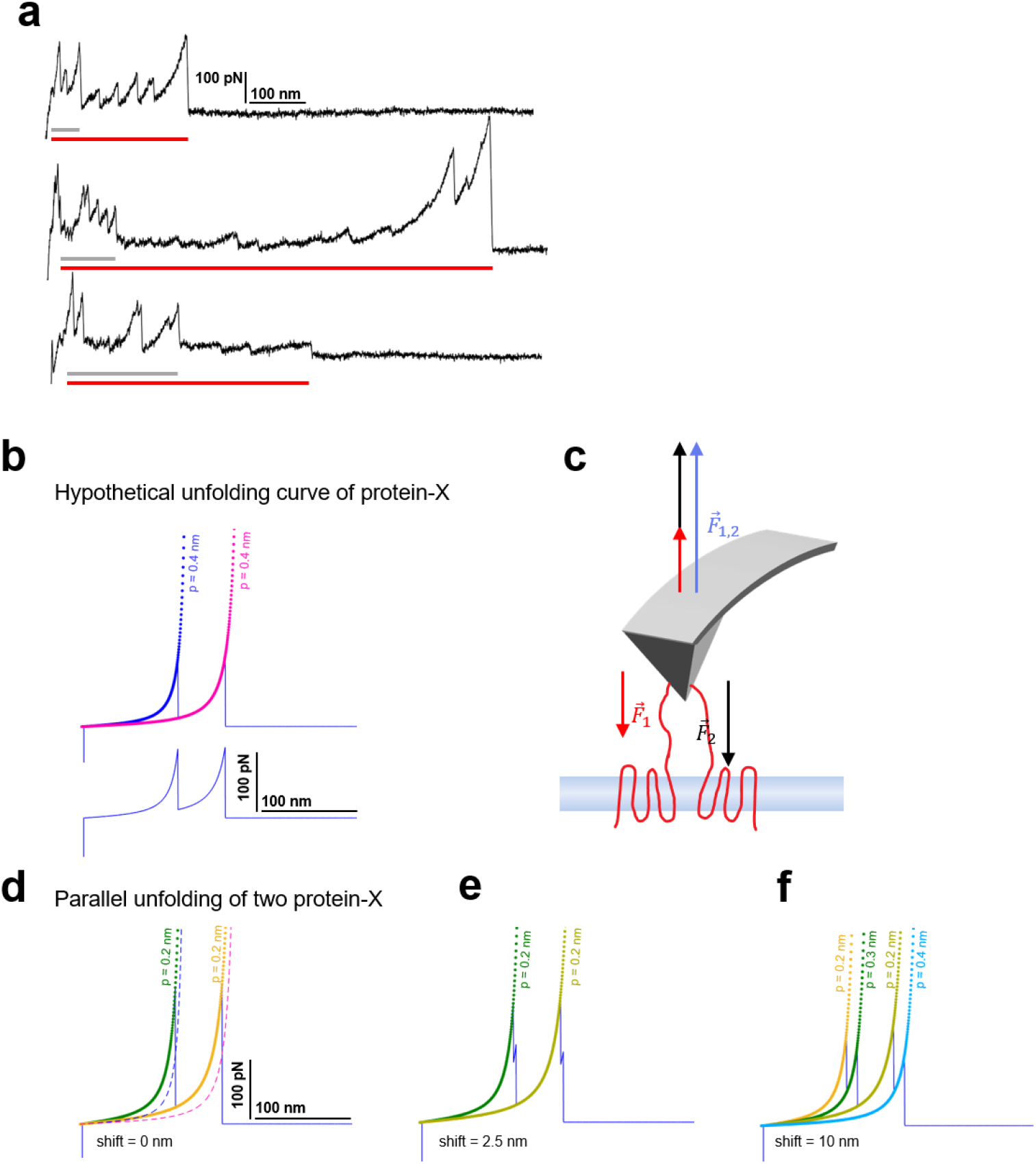
Candidates of multiple unfolding and origin of persistence length deviation. **a,** observed F-d curves showing the features of multiple unfolding events (red horizontal bar: long protein, gray horizontal bar: short protein; guessed). **b**, hypothetical unfolding curve of protein-X (peak 1: Lc=100 nm, F=150pN; peak 2: Lc=150 nm, F=150pN) fitted with the WLC model with standard persistence length p=0.4nm. **c**, the force applied by the AFM tip balances the unfolding forces of the two proteins during the retraction. **d**, the effective F-d curve recorded during parallel unfolding of two protein-X corresponds to the sum of a single unfolding curve **b** and it is best fitted with half the persistence length p=0.2nm. **e**, relative shift of 2.5 nm and **f**, 10 nm still result in deviations of the measured persistence length and display the doubling of the peaks.

**Figure S8.**
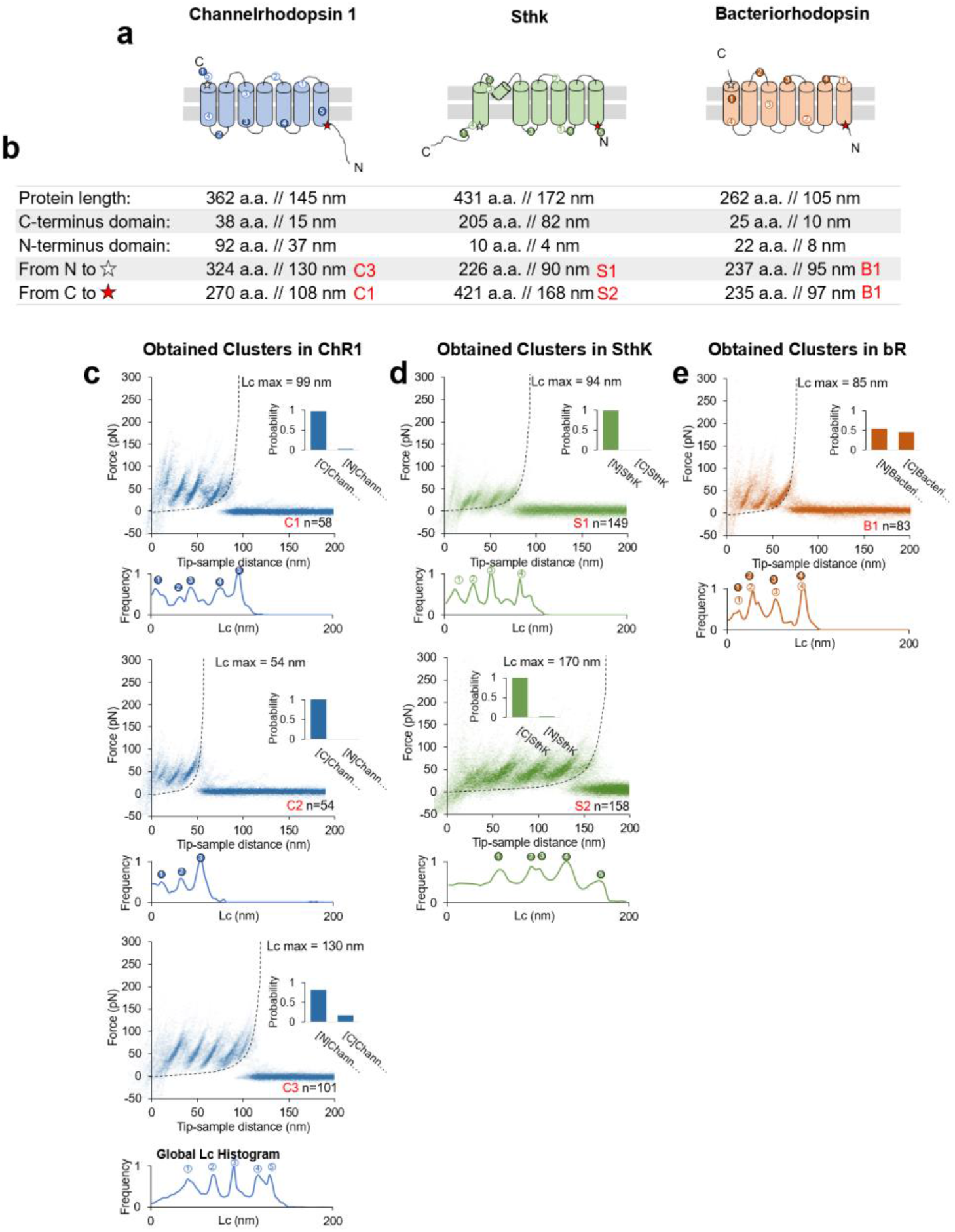
Unfolding of reconstituted membrane proteins. **a**: scheme of the structure of the three purified proteins used in the in vitro preliminary step: ChR1, Sthk and bR (cylinders represent alpha helices). **b**: geometrical length of the amino-acid chains constituting the whole protein, the C and N-terminals and of the length of the whole protein minus the C and N-terminal. The codes in red represent the name un the respective unfolding cluster. **c**: clusters obtained from a dataset of n=18868 traces from a sample with ChR1. The histogram bars on the right of each density plot represent the Bayesian assignment according to our procedure (described in Methods and in Fig. 6). [N] and [C] stand for “protein unfolded from N-terminus/C-terminus”. Below each density plot the corresponding Global histogram of contour length showing the most likely position for the unfolding peaks to occur: the circles with numbers are then reported in **a** to show the position of the preferential unfolding position in the structure. **d**: clusters obtained from a dataset of n=115155 traces from a sample with SthK. **e**: clusters obtained from a dataset of n=35599 traces from a sample with bR.

**Figure S9.**
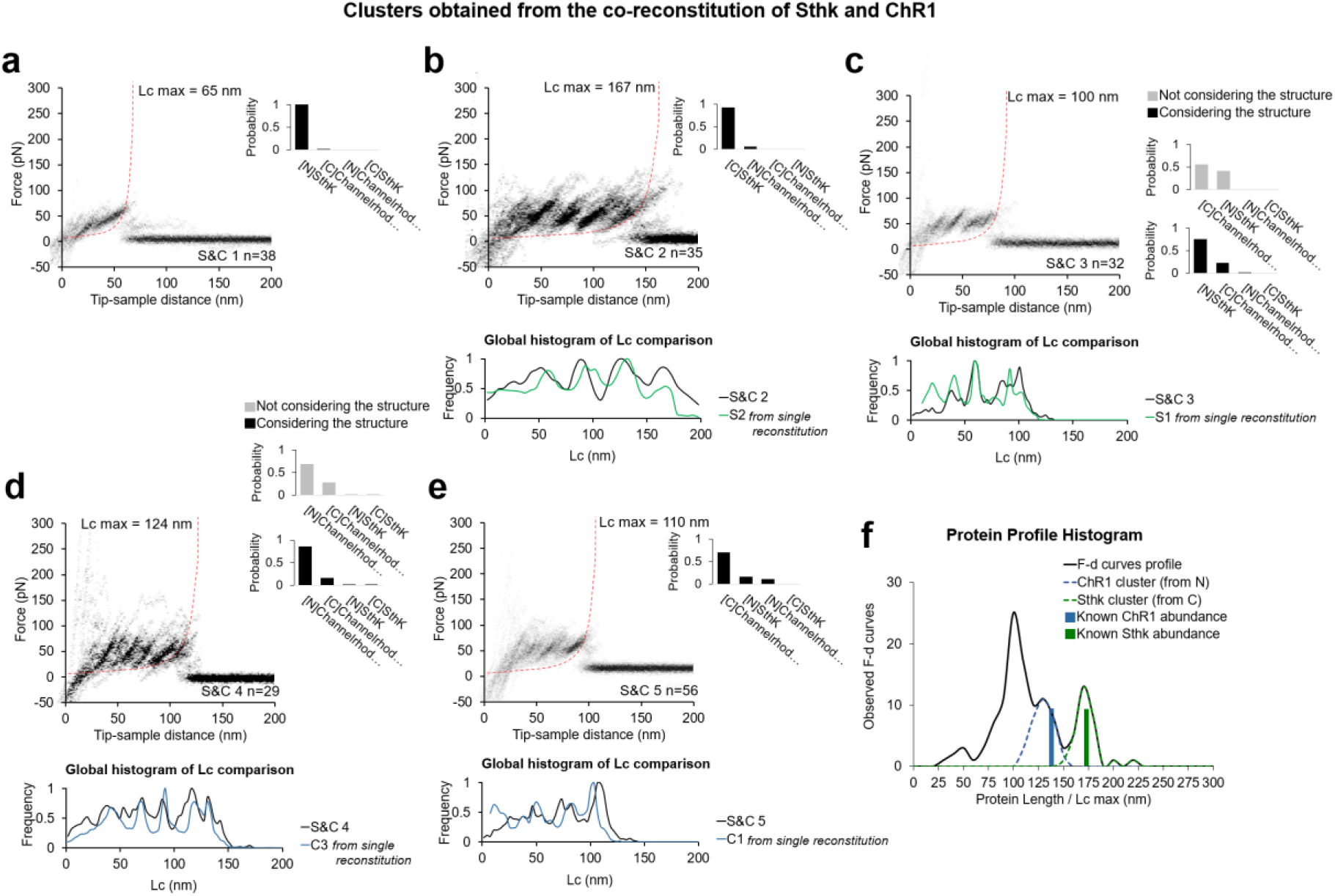
In vitro experiments with co-reconstitution of SthK and ChR1 (relative concentration 1:1) showing the five clusters obtained, and comparison of these new patterns S&Cx to the previous Cx and Sx clusters from single reconstitution. **a-e** Clusters (Called S&C 1, …, S&C 5) resulting from the simultaneous reconstitution of ChR1 and SthK in a concentration ratio of 1:1, and Global histogram of the contour length comparison with the best candidate obtained from the single reconstitution (from Fig. 8). Bayesian probability shown on the right of each cluster. For **c** and **d** we report the improvement in the Bayesian identification when the structure of the protein is taken into account (see methods Bayesian estimation). **f:** black line: distribution of the maximum of the contour length (Lc max) of F-d curves of the clusters (except **a**) obtained in the co-reconstituted sample; blue and green bars: real length of the protein in the sample.

**Figure S10.**
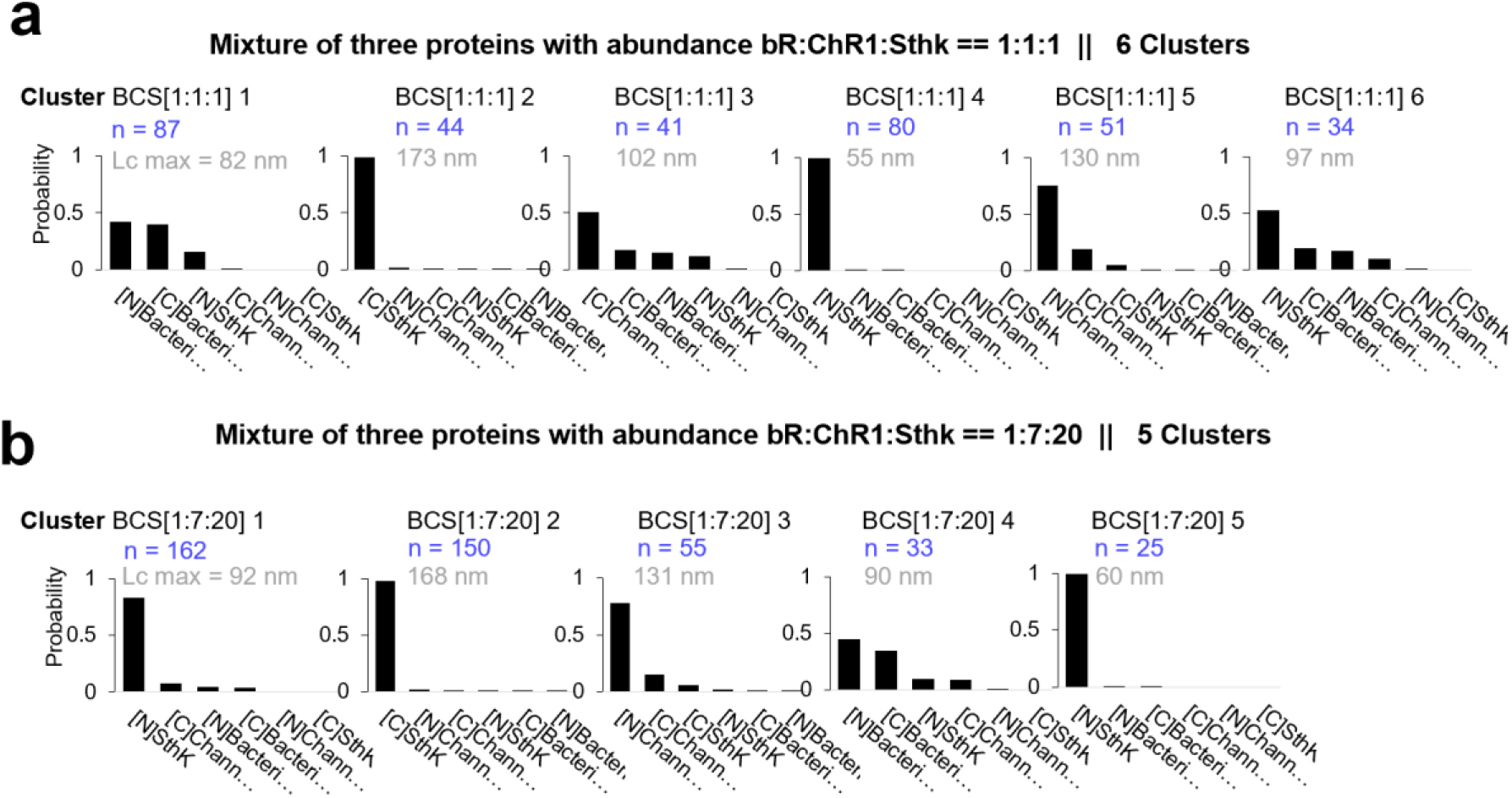
Clusters obtained in *in vitro* experiments with mixtures of bR, ChR1 and Sthk at different relative concentrations [1:1:1] and [1:7:20]. Cluster names: BCS[1:1:1]x and BCS[1:7:20]x. **a:** candidates’ prediction of the clusters obtained in samples with bR, ChR1 and Sthk and relative abundance of [1:1:1]. (The number in gray indicates the total contour length of the clusters called Lc max, the number in blue the number of traces per cluster). **b:** same as **a** for relative abundance of [1:7:20].

**Figure S11.**
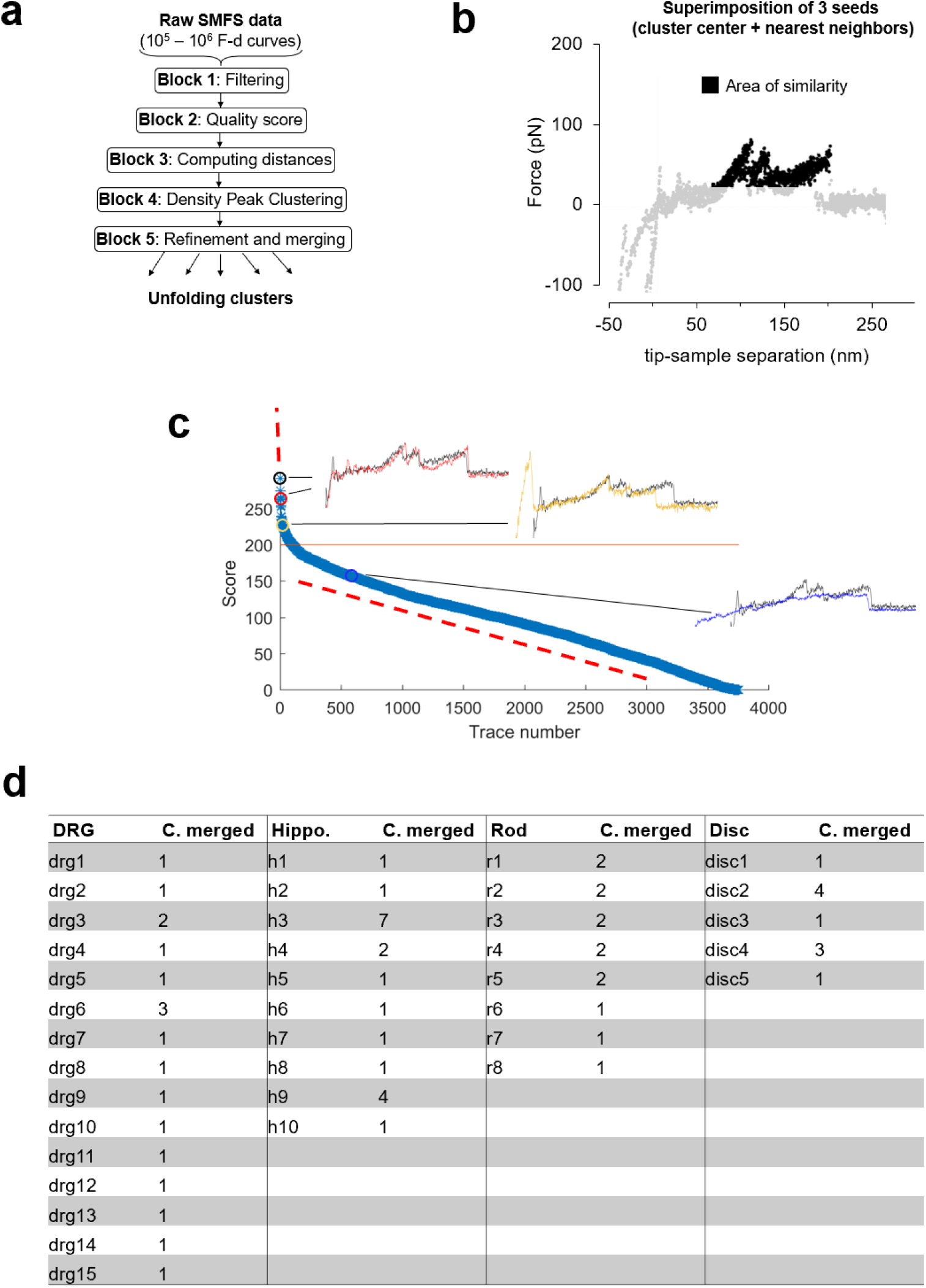
Clustering. **a**, block scheme of the clustering method. **b**, area of similarity (AoS) for cluster R1 used for block 5 described in Methods. **c**, plot of the scores generated by the AoS on **b** of the F-d curves in the Dataset Rod filtered by length in the descending order. **d**, table showing the number of clusters that was merged to form the final selection of Fig. 3.

**Figure S12.**
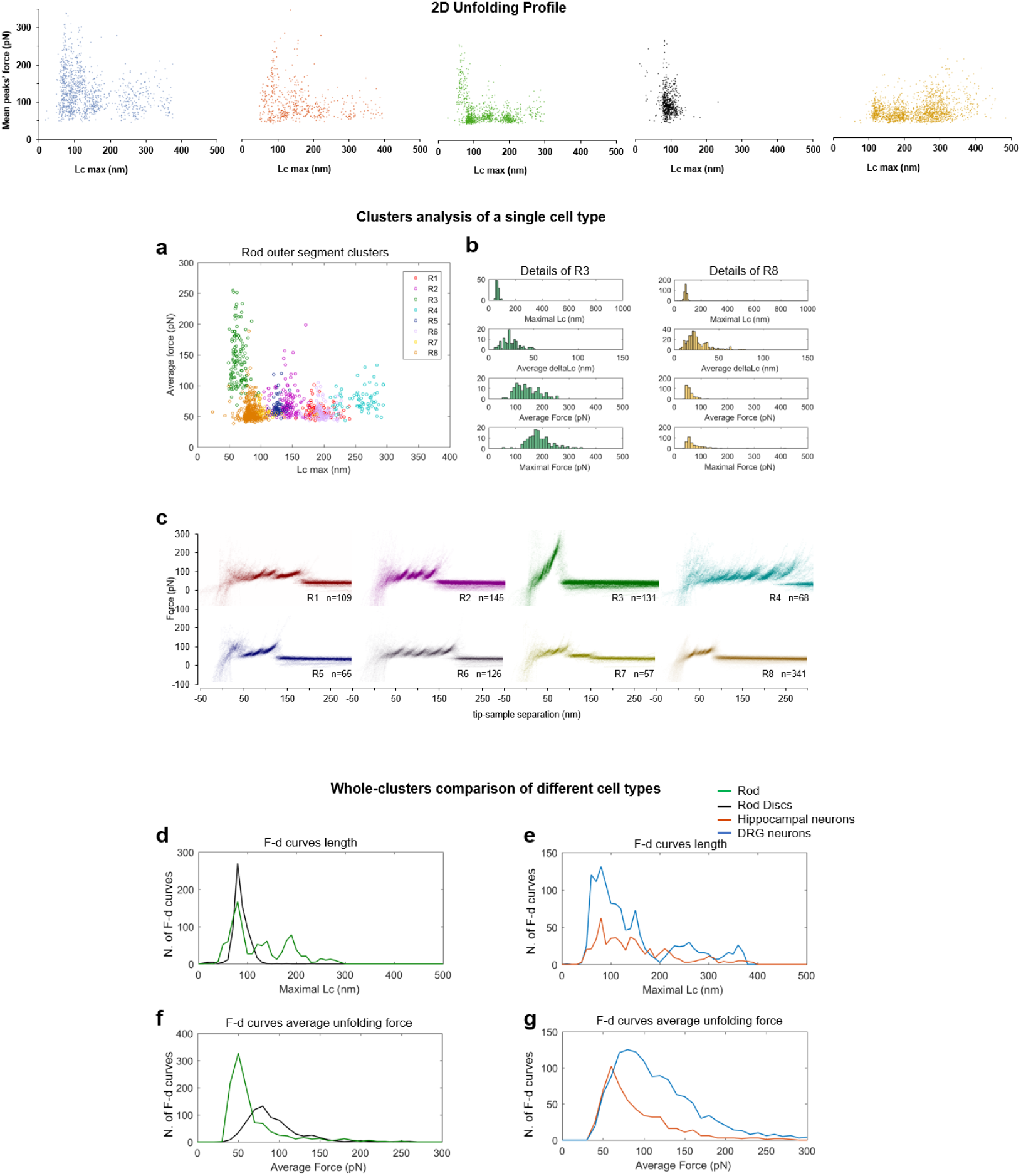
Alternative visualizations of cluster analysis. **Top,** two dimensional unfolding profile in the compact representation of all the clustered F-D curves: x-axis is the maximum contour length; y-axis is the average unfolding force (DRG: n = 1255; hippocampus: n = 563; rod: n = 1039; disc: n = 703, NG108: n = 1591)**. a**, plot of all the F-d curves belonging to the clusters of the rod outer segment plotted with different colors in the ‘average unfolding force’ vs ‘maximal contour length’ space. **b**, distribution of representative observables for clusters R3 and R8. **c**, density plots of clusters in **a**. Comparison of the maximal contour length profiles (**d** and **e**) and of the average unfolding forces (**f** and **g**) of the four cell types investigated.

**Figure S13.**
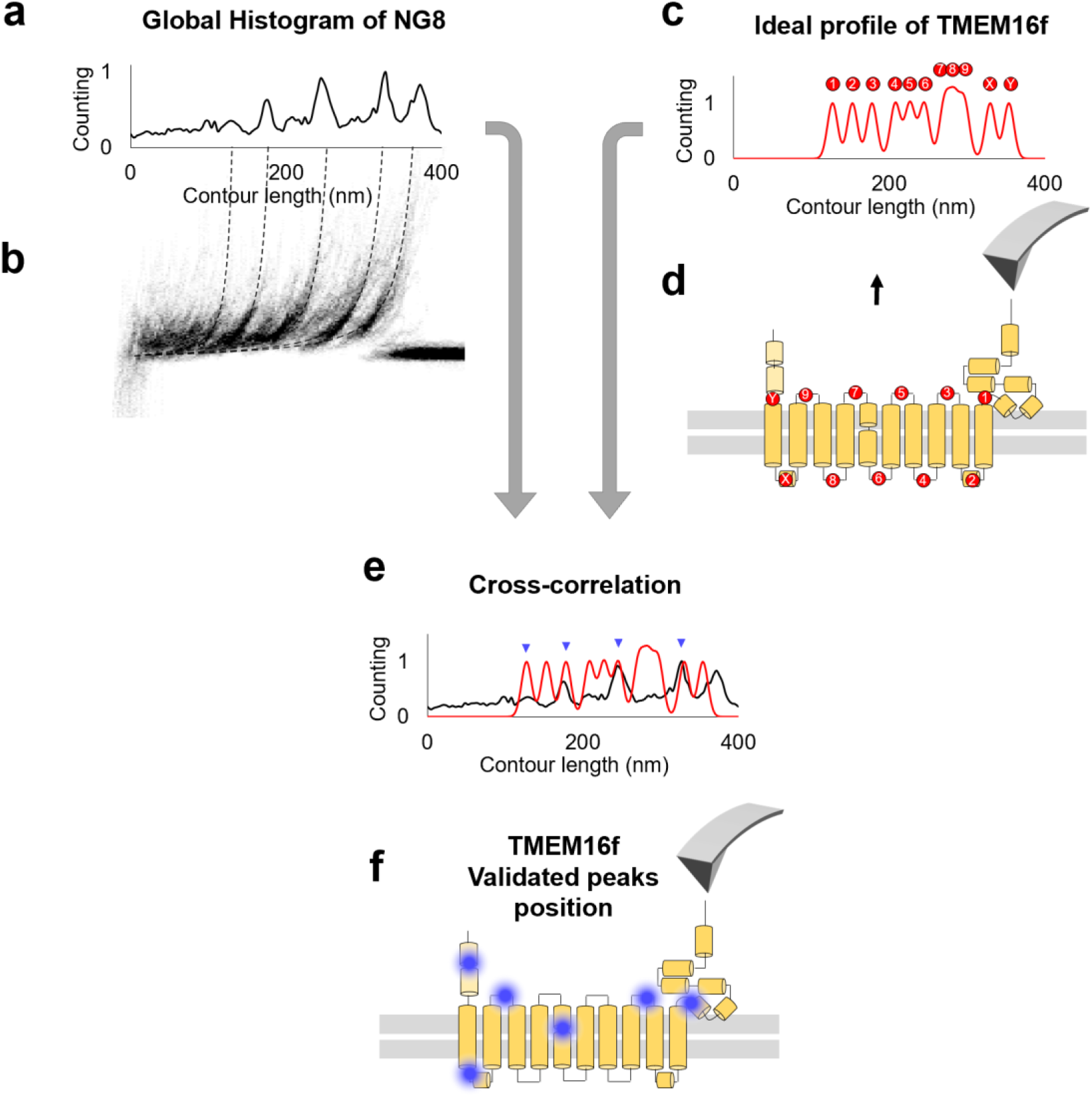
**The cross-correlation used to evaluate** P(Peaks_Cx_|Structure_ProtA_). **a**: Global Histogram of cluster NG8 and the density plot (**b**). The peaks in **a** represent the region where the peaks tend to occur. **c**: ideal profile of TMEM16F built starting from the information that 86% of the peaks happen in the loops (**d**). **e:** superimposition of the real and the idealized profile: the correlation between the two generates a number that is used in the probability estimation. The more peaks correspond to loops, the higher is the cross-correlation value. **f**: Blue dots indicate the location of force peaks in the Cryo-EM structure of the TMEM16F.

**Figure S14.**
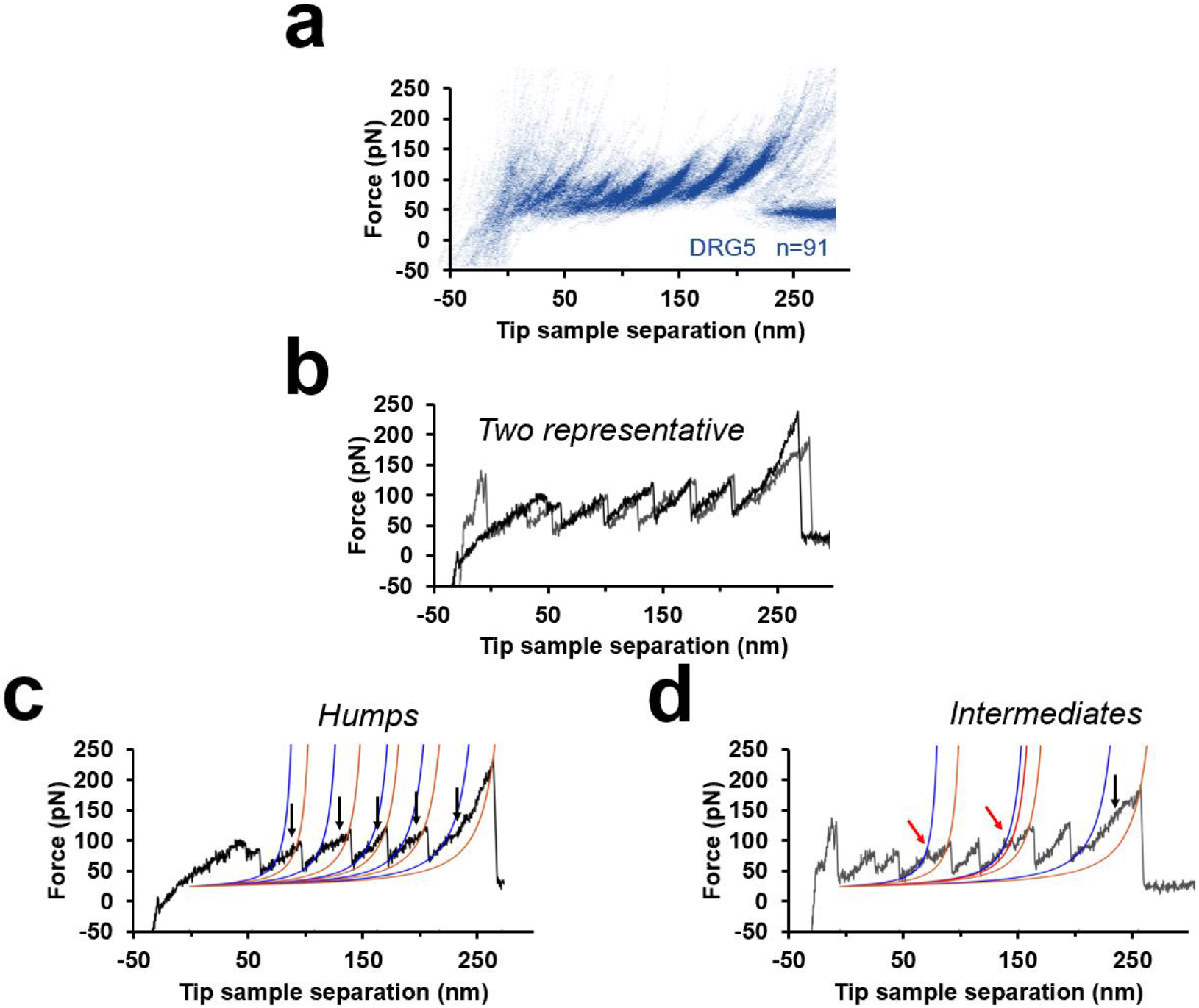
Unfolding intermediates and humps a, cluster DRG5 from Fig. 3. b, two representative F-D curves from DRG5 (black and gray). c, Black curve in b that shows humps (blue WLC fit identify the beginning of the humps, gold WLC fit identify the end). d, gray curve in b that shows multiple peaks (red arrows) instead of humps.

**Figure S15.**
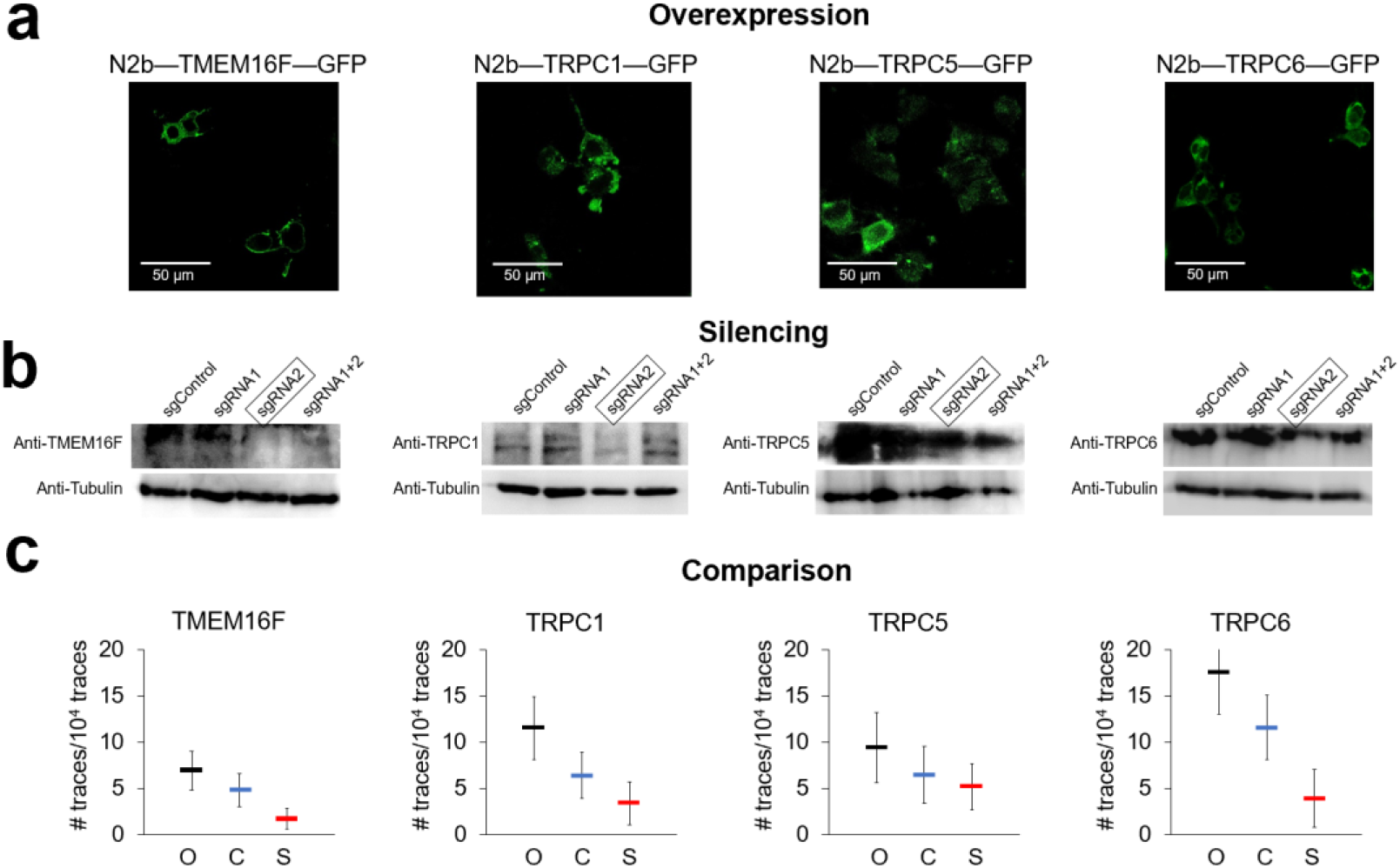
Overexpression and silencing in NG108-15 cells. **a:** fluorescent confocal images of the cells overexpressing the 6xHis-N2B—Protein—GFP construct, where Protein was TMEM16F, TRPC1, TRPC5 and TRPC6. **b**: Western Blots for the four proteins indicating the silencing obtained with different plasmids (the chosen plasmids for the SMFS experiments are the highlighted ones). **c**: For each cluster of F-d traces corresponding to TMEM16F, TRPC1, TRPC5 and TRPC6 we considered the frequency of identified traces over 10000 F-d traces which passed our criteria of good/valid traces associated to the unfolding of a membrane protein. (C) Control F-d traces were those obtained from not transfected cells. (O) Overexpressed F-d traces were those obtained from cells transfected with the construct 6xHis-N2B-Protein-GFP bearing the N2B signature and those without the N2B signature but classified as corresponding as the unfolding of the Protein under investigation. (S) Silenced F-d traces were those obtained from NG108-15 cells in which the gene coding for the protein under investigation was silenced. Error bars indicate the standard deviation.

**Table S1.**
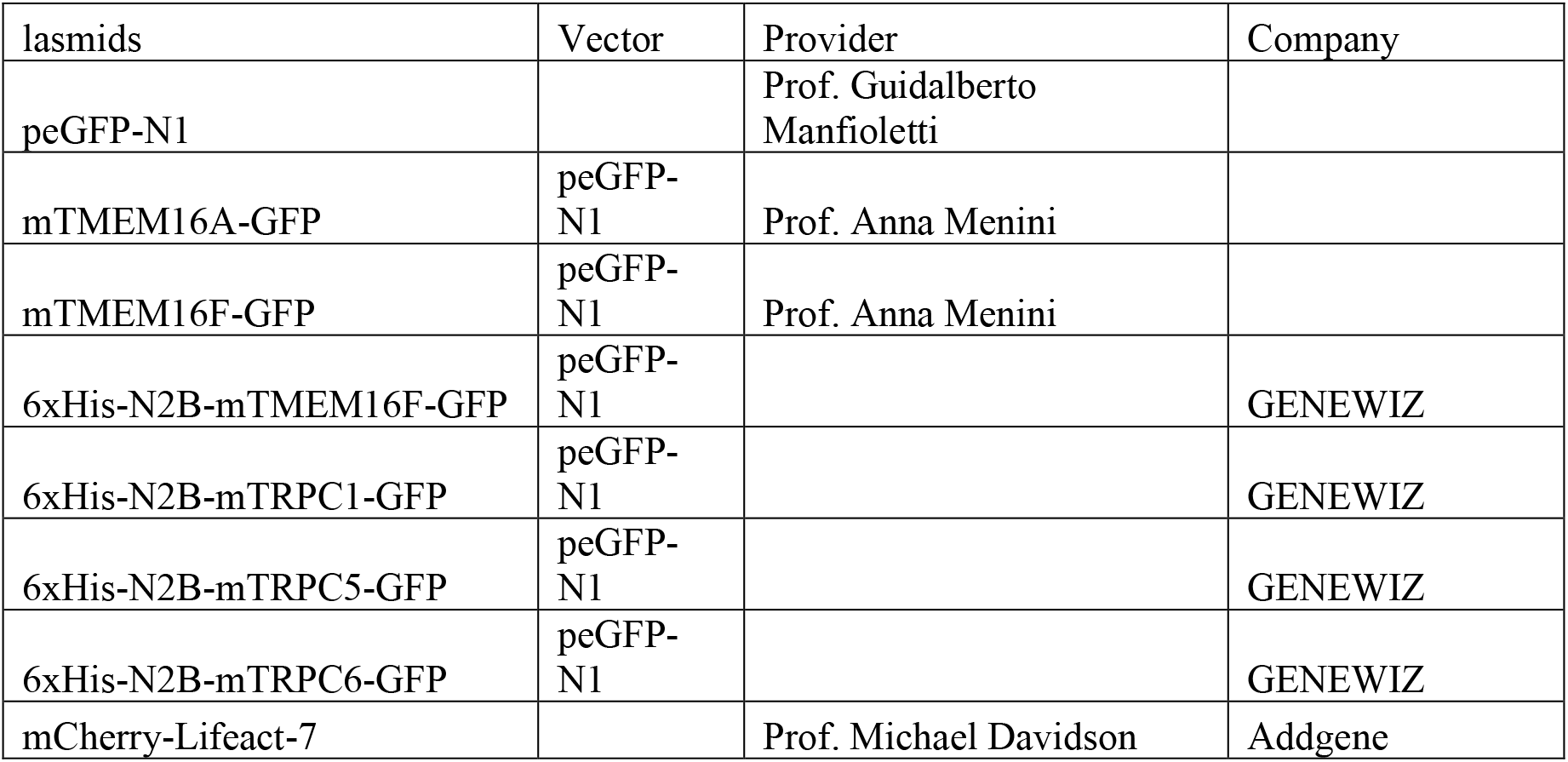
List of plasmids. The cDNA of TRPC1/5/6 were from Miao Ling Plasmid Company. All the 6xHis-N2B-Protein-GFP construct is made by Genewiz company.

**Table S2.**
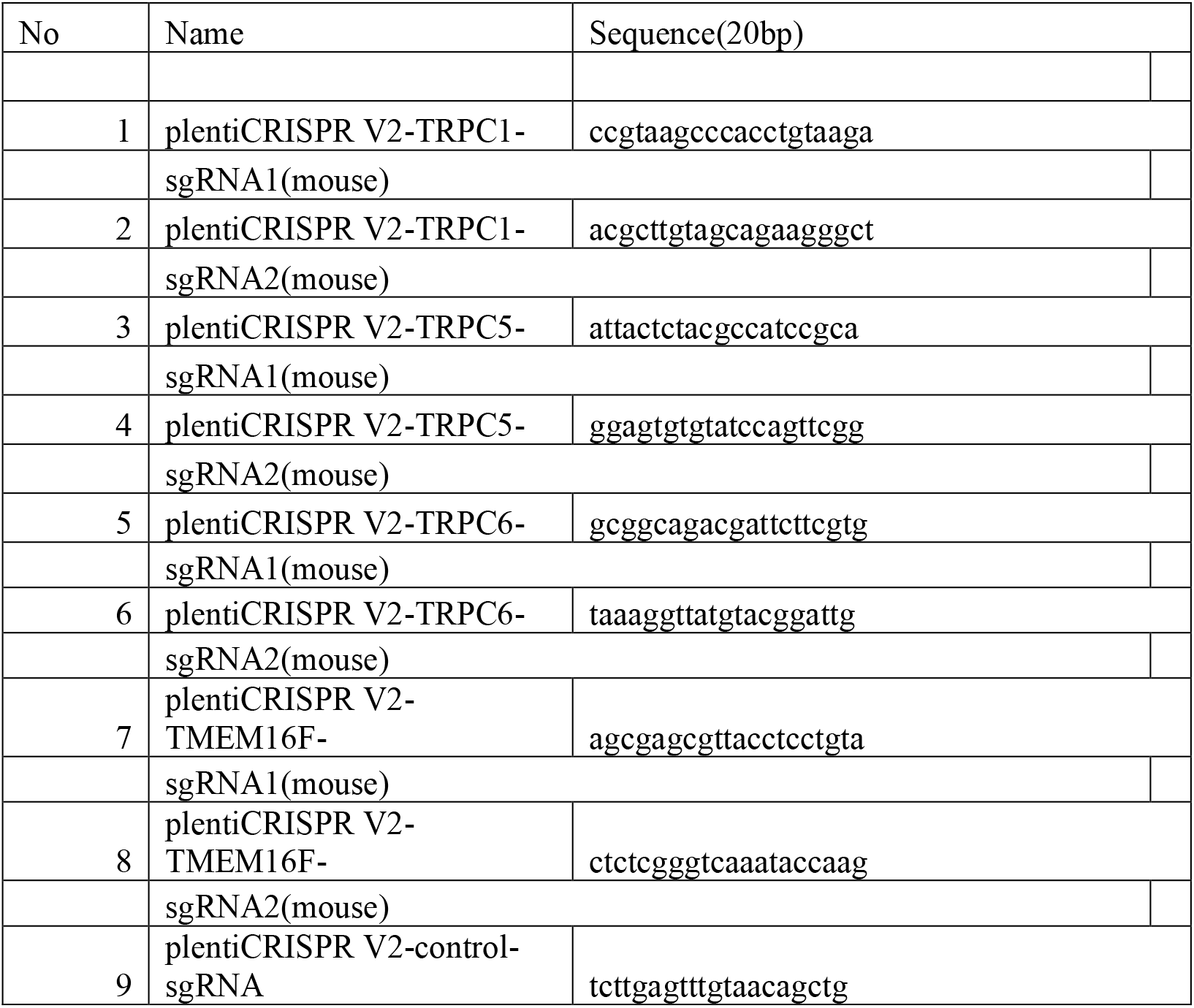
sgRNA Sequences. The sgRNA sequences cloned into LentiCRISPR-V2 puro vector were purchased from Miao Ling Plasmid Company.

**Data S1. (separate file)**

Tables with the protein lists and variables used for the Bayesian identification.

